# Co-aggregation with Apolipoprotein E modulates the function of Amyloid-β in Alzheimer’s disease

**DOI:** 10.1101/2021.07.13.452239

**Authors:** Zengjie Xia, Emily E. Prescott, Hollie E Wareing, Martyna M Matuszyk, Helen Dakin, Eleni Dimou, Eric Zuo, Yu P. Zhang, Jeff Y.L. Lam, John S. H. Danial, Tom Leah, Katy A. Barnes, Michael Strickland, Hong Jiang, Peter Thornton, Damian C. Crowther, David M. Holtzman, Simon M. Bell, Adrian Higginbottom, Laura Ferraiuolo, Heather Mortiboys, Stephen B. Wharton, Rohan T. Ranasinghe, David Klenerman, Suman De

## Abstract

Which isoforms of apolipoprotein E (apoE) we inherit determine our risk of developing late-onset Alzheimer’s Disease (AD), but the mechanism underlying this link is poorly understood. In particular, the relevance of direct interactions between apoE and amyloid-β (Aβ) remains controversial. Here, single-molecule imaging shows that all isoforms of apoE associate with Aβ in the early stages of aggregation and then fall away as fibrillation happens. ApoE-Aβ co-aggregates account for ∼50% of the mass of soluble Aβ aggregates detected in the frontal cortices of homozygotes with the higher-risk *APOE4* gene. Our results connect inherited *APOE* genotype with the risk of developing AD by demonstrating how, in an isoform- and lipidation-specific way, apoE modulates the aggregation, clearance and toxicity of Aβ. Selectively removing non-lipidated apoE4-Aβ co-aggregates enhances clearance of toxic Aβ by glial cells, and reduces inflammation and membrane damage, demonstrating a clear path to AD therapeutics.

## Introduction

Inherited variation in the sequence of apolipoprotein E (apoE) is the greatest genetic risk factor for late-onset Alzheimer’s Disease (AD), which accounts for ∼95% of all AD cases^1^. The *APOE* gene has three alleles: *APOE2, APOE3*, and *APOE4*. Compared to the most common *APOE3* form, *APOE2* is neuroprotective^2^, while *APOE4* increases AD risk; *APOE4* homozygotes are 15-fold more susceptible to AD than *APOE3* homozygotes^3,4^, and disease begins several years earlier in *APOE4* carriers than those with *APOE3* or *APOE2*^3^.

Attempts to establish how apoE influences AD risk have focused on its effect on the pathological amyloid-beta (Aβ) peptide^1,5^. Deposits of Aβ aggregates in the central nervous system (CNS) are a hallmark of AD^6^, and disturbed Aβ homeostasis is one of the earliest events in AD pathogenesis^7,8^. This dysfunction induces synaptic and axonal damage as well as tau seeding and spreading, leading to neurodegeneration in AD^9–11^. There is compelling evidence that apoE4 enhances Aβ pathology^5^: carriers of the *APOE4* gene have more Aβ deposits in their CNS than non-carriers^1,12^, exhibit amyloid positivity earlier in life^13^, and experience a faster-growing Aβ burden^14^. The isoforms of the apoE protein seem to differentially affect the levels of Aβ in the CNS, which may explain their differential AD risk profiles. Lipidation of apoE may also play a role in AD. Although mostly lipidated in the human CNS^5,15^, poorly or non-lipidated apoE increases Aβ pathology^16^, and the risk of developing AD^17^. Observational studies show that apoE influences the relationship between Aβ and cognitive decline in AD^18–20^. However, it is unclear how the different isoforms and lipidation states of apoE affect Aβ aggregation, clearance, and aggregate-induced neurotoxicity at the molecular level.

The faster Aβ plaque deposition in carriers of *APOE4*^14,21^ has led to several hypotheses for apoE4’s molecular role in AD. One possibility is that apoE4 promotes more aggregation of Aβ than the other isoforms by interacting directly with Aβ when the two meet in the extracellular space. The discovery of co-deposited apoE in AD amyloid plaques provided early circumstantial evidence for this idea^15,22,23^, but biophysical studies are equivocal: apoE can either speed up or slow down Aβ aggregation *in vitro*, depending on the conditions^24^. Targeting non-lipidated apoE in amyloid plaques with an antibody can significantly reduce Aβ pathology in transgenic mouse models^15,22^ as well as reduce Aβ-induced tau seeding and spreading^25^. However, studies disagree on whether soluble forms of apoE and Aβ associate significantly in brain tissue^17,26^. The role of apoE-Aβ interactions in Aβ clearance is also unclear: while apoE might traffic Aβ out of the interstitial brain fluid^21,27^, other work suggests that apoE and Aβ instead compete for clearance-mediating receptors^26^.

In this study, we used single-molecule imaging to monitor how apoE affects the form, function, and clearance of Aβ aggregates at different stages of maturity. We show that apoE influences the structure and composition of Aβ aggregates during oligomerization, and that apoE-Aβ interactions tune disease-relevant functions of Aβ in an isoform- and lipidation-specific manner. Our work identifies a role for transient, non-lipidated apoE-Aβ co-aggregates in modulating Aβ deposition and neurotoxicity, and thereby connects *APOE* genotype inheritance with the risk of developing sporadic AD.

## Results

### Soluble apoE-Aβ Co-aggregates Form Isoform-independently, but Accumulate Isoform-dependently in AD Brains

We began our study by investigating how apoE interacts with Aβ along its aggregation pathway to fibrils. Aggregating Aβ comprises a dynamic, heterogeneous mixture of species with different sizes, shapes, and properties, and small sub-populations may disproportionately contribute to AD^28,29^. We reasoned that apoE could influence AD risk by interacting with Aβ aggregates that are transient and/or rare, which might explain why previous attempts to study association by taking snapshots of the bulk mixture have painted an inconsistent picture. We therefore aggregated Aβ42 *in vitro* at a concentration (4 µM) that would give rise to fibrils, in the presence and absence of near-physiological concentrations of each non-lipidated and lipidated apoE isoform (∼80 nM)^30,31^.

Assaying fibril formation using ThT fluorescence confirmed that all reactions produced fibrils and that the presence of non-lipidated apoE slowed aggregation, while the presence of lipidated apoE sped the reaction up (Figure 1A, Figure S1); lipid particles are known to accelerate the aggregation of amyloidogenic proteins by providing a surface for primary nucleation^32–34^. In order to sample the heterogeneity within each aggregation pathway, we characterized individual aggregates at different stages of the reaction. We imaged each aggregation reaction at the end of the lag phase (t_1_), the middle of the growth phase (t_2_), and the plateau phase (t_3_), using single-molecule pull-down (SiMPull)^35^ (Figure 1B). In this assay (Figure 1C), Aβ42 is captured using a surface-tethered 6E10 antibody and imaged using two-color total internal reflection fluorescence (TIRF) after adding primary detector antibodies for Aβ (Alexa-Fluor-647-labeled 6E10) and apoE (Alexa-Fluor-488-labeled EPR19392) (Figure S2). Using the same monoclonal antibody to sandwich Aβ aggregates renders unreacted monomers undetectable because they only contain one epitope.

**Figure 1.**
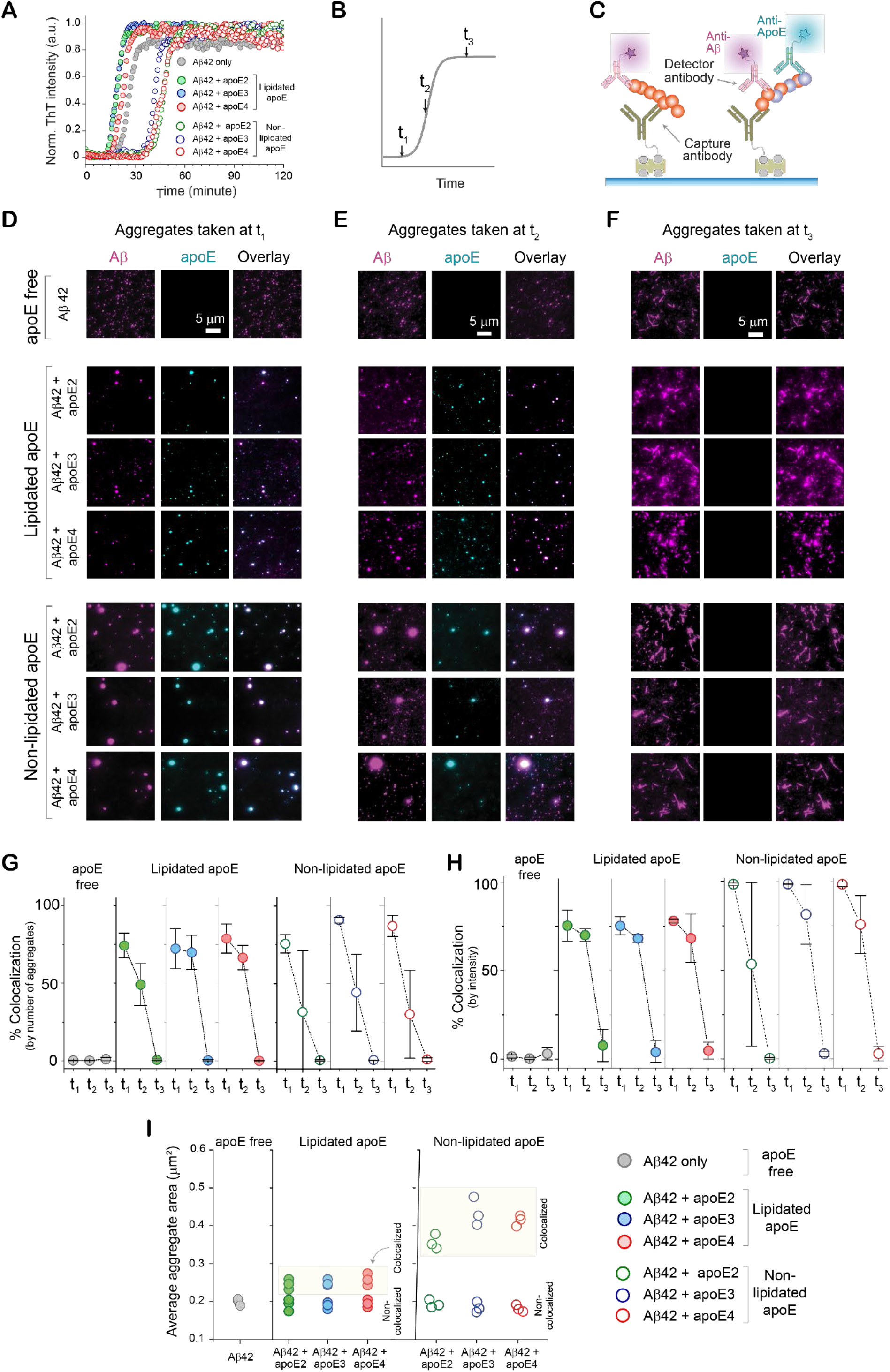
ApoE and Aβ transiently co-aggregate en route to fibrils. **(A)** Aβ42 aggregation (4 µM) in the presence of different non-lipidated isoforms of apoE (0 or 80 nM), monitored by ThT fluorescence (*n* = 3 independent replicates). **(B)** Time points at which samples were taken for further analysis. **(C)** SiMPull assay for Aβ42 aggregates and apoE-Aβ co-aggregates (1 µM Aβ monomer equivalents), using biotinylated 6E10 antibody for capture, and Alexa-Fluor-647-labeled 6E10 (500 pM) and Alexa-Fluor-488-labeled EPR19392 (1 nM) antibodies for detection. **(D-F)** Two-color TIRF images of aggregates were captured at t_1_ **(D)**, t_2_ **(E)**, and t_3_ **(F). (G-H)** Colocalization between Aβ and apoE at different time points, quantified by aggregate counting **(G)** and 6E10 fluorescence intensity **(H)**. Data are plotted as the mean and standard deviation of three technical replicates. **(I)** Average sizes of colocalized and non-colocalized aggregates (N.B. in diffracted-limited imaging, the minimum apparent aggregate size is ∼0.2 µm^2^).

Characterizing individual aggregates rather than their ensemble average allowed us to extract properties of the heterogeneous population including size, shape, and composition; aggregates containing apoE and Aβ42 should be colocalized in both detection channels. These images (Figure 1D-F) revealed that all types of apoE coaggregate with Aβ42 in the early stages (t_1_ and t_2_) of aggregation, but that co-aggregates disappear as the reaction reaches completion. There was no significant isoform dependence in the extent of colocalization, whether based on the aggregate number (Figure 1G), or intensity (Figure 1H). These findings are independent of the antibody used to detect apoE (Figure S3), and there is no colocalization when isotype-control detection antibodies are used (Figures S4 and S5), suggesting minimal contributions from non-specific binding. Using the intensity as a proxy for the amount of protein present in each aggregate suggests that ∼75-100% of aggregate mass at the end of the lag phase is in co-aggregates, which falls to ∼30%-60% in the growth phase, and ∼0% by the plateau phase. Although non-lipidated apoE colocalizes with Aβ42 at slightly higher levels, the main effect of lipidation was on the apparent size of co-aggregates (Figure 1I): those containing non-lipidated apoE (mean diameter ∼500-900 nm) were much larger than those containing lipidated apoE (mean diameter ∼200-250 nm or less; in diffraction-limited imaging, any aggregates smaller than the diffraction limit will appear in this range).

Our finding that non-lipidated apoE forms large soluble co-aggregates with Aβ supports the idea that apoE in this form can stabilize soluble species formed early on in aggregation and thus inhibit Aβ fibrillation^36^. Because we used apoE in sub-stoichiometric amounts (Aβ: apoE 50:1), it is unlikely to slow down aggregation by sequestering Aβ monomers. ApoE more likely interacts with Aβ42 in its earliest stages of aggregation; apoE did not associate with preformed Aβ42 aggregates (Figure S6). The high fluorescence intensities of these soluble co-aggregates indicate high effective protein concentrations, which may seem incompatible with decreased fibrillization rates. However, it has recently been shown that locally concentrating Aβ42 in condensates can significantly slow its aggregation^37^. The fact that fibrils have shed all associated apoE might indicate that elongating heteronuclei is less energetically favorable than elongating homonuclei. Importantly, the lack of any isoform dependence means that co-aggregation alone cannot explain the *APOE* dependence of AD.

To determine whether these apoE-Aβ co-aggregates are disease-relevant, we isolated Aβ aggregates from the postmortem frontal cortices of six AD patients (Supplementary Table 1). We chose a method that gently extracts soluble aggregates by soaking^38^, in preference to tissue homogenization, which extracts insoluble, predominantly fibrillar aggregates that are mostly inert^39^. We imaged the extracts from three homozygous *APOE4* and three homozygous *APOE3* AD patients using SiMPull (Figure 2A). All samples yielded Aβ aggregates both with and without associated apoE; extracts from *APOE4* carriers contained more of both types of aggregates than extracts from *APOE3* carriers (Figure 2B). This finding supports the previous report that *APOE4* carriers have more soluble Aβ aggregates than *APOE3* carriers^40,41^. Unlike our *in vitro* aggregations - where the initiation of aggregation is synchronized for all monomers - only ∼5% of Aβ aggregates in *APOE4* homozygotes and ∼1% of aggregates in *APOE3* homozygotes are co-aggregates (Figure 2C). This difference probably results from the transience of co-aggregates and the fact that Aβ and apoE are replenished and cleared over years in the CNS. Extracted aggregates therefore reflect a broad sample of the reaction pathway, *e*.*g*. recently formed soluble aggregates alongside mature fibrils. Although few in number, some of the *ex vivo* co-aggregates are large (Figure 2E) and contribute disproportionately to the fluorescence intensity. Using the intensity as a proxy for total amounts of protein suggests that co-aggregates comprise 40-60% and 10-35% of the Aβ-aggregate mass in *APOE4* and *APOE3* carriers, respectively (Figure 2D). These data, therefore, reveal a large isoform dependence in the accumulation of co-aggregates in AD brains.

**Figure 2.**
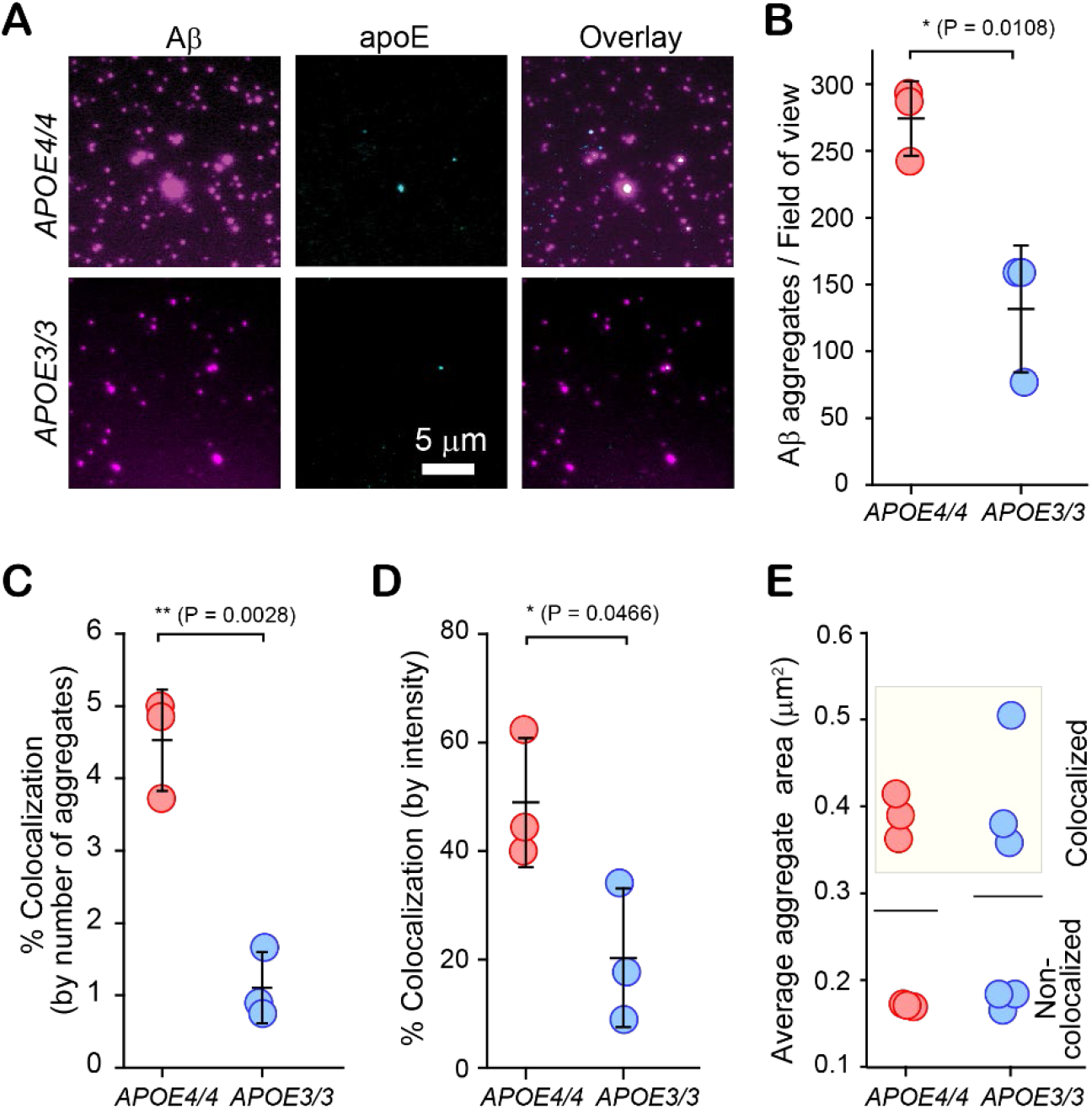
Aβ-apoE co-aggregates form in human brain tissue, but their concentration is isoform-dependent. **(A)** Two-color TIRF images of aggregates from frontal-cortex extracts of homozygous *APOE4* and *APOE3* AD patients. **(B)** Numbers of Aβ-containing species captured from *APOE4* and *APOE3* homozygotes. **(C, D)** Colocalization between Aβ and apoE in extracts from *APOE4* and *APOE3* homozygotes, quantified by aggregate counting **(C)** and 6E10 (Aβ) fluorescence intensity **(D). (E)** Sizes of apoE-colocalized and non-colocalized Aβ aggregates in *APOE4* and *APOE3* homozygotes. Data points in panels **B-E** represent one of three biological replicates; error bars represent standard deviation. Statistical significance was calculated using a two-sample t-test. *P < 0.05, **P < 0.01.

### Glial Uptake of Soluble apoE-Aβ Co-aggregates is Isoform- and Lipidation-dependent

Given that all isoforms of apoE, whether lipidated or not, had co-aggregated to similar extents with Aβ42, we next asked whether differential clearance might explain the isoform-dependent accumulation of apoE-Aβ co-aggregates in AD brains. We quantified the uptake of aggregates by two types of cells: i) immortalized murine microglial BV-2 cells, and ii) astrocytes isolated from the cerebral cortex of humans (Figure 3A-C). BV-2 cells and human astrocytes take up soluble and insoluble Aβ aggregates^42,43^, and dysfunction of microglia and astrocytes is implicated in AD^44,45^. Alongside monitoring aggregate clearance by these cells, we also measured the resulting neuroinflammation. Protein aggregates cause neuroinflammation by disrupting CNS homeostasis^46^, and neurotoxic Aβ aggregates are known to promote pro-inflammatory pathways in microglia^44^ and astrocytes^45^.

**Figure 3.**
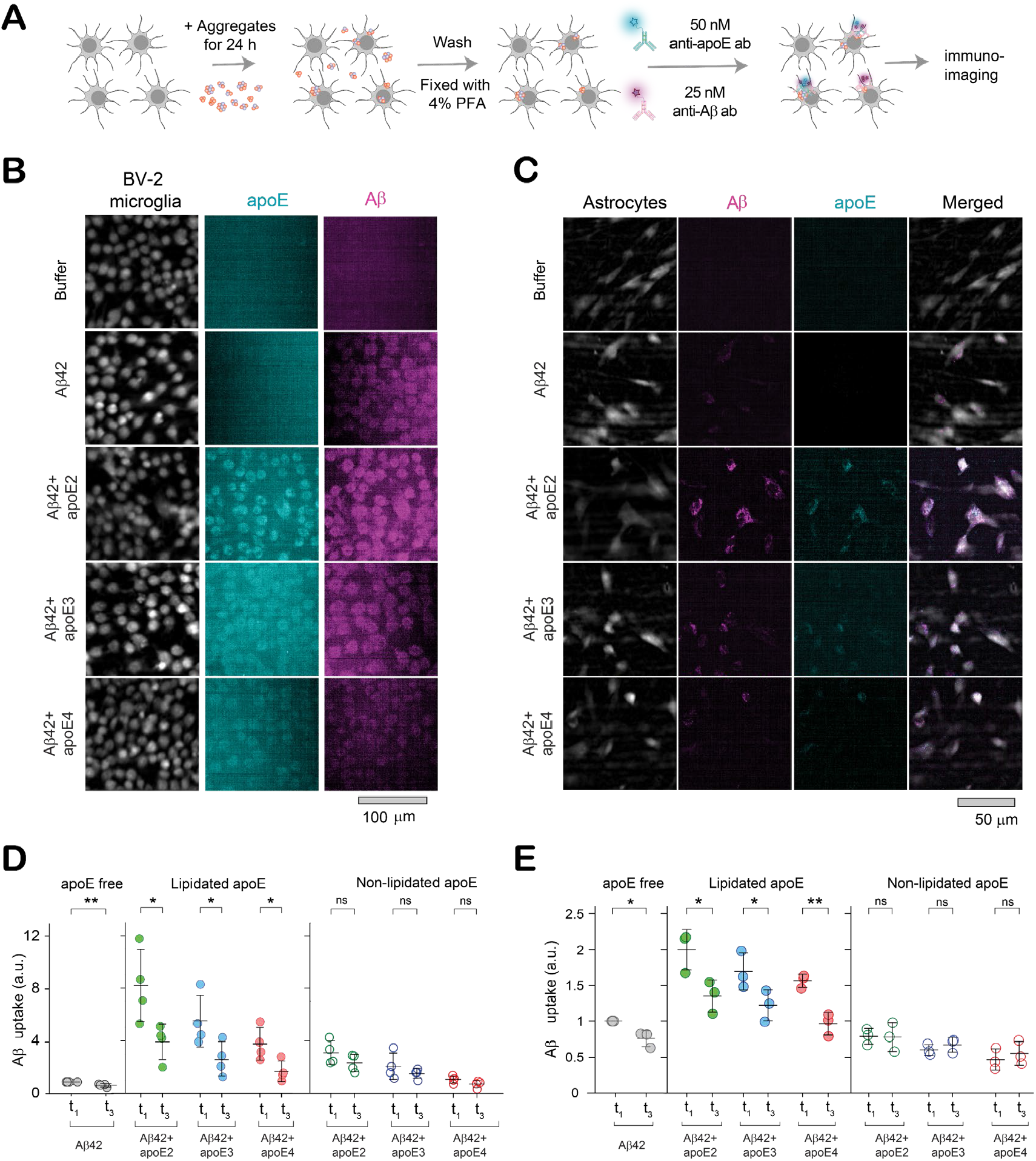
Clearance of apoE-Aβ co-aggregates by glial cells depends on aggregate maturity, apoE isoform and lipidation. **(A)** Assay for uptake of aggregates by glial cells. **(B-C)** Two-color epifluorescence images showing uptake of Aβ42 (4 µM monomer equivalents) and apoE (0 or 80 nM) from aggregation mixtures at t_1_ (end of lag phase) and t_3_ (plateau phase) by BV-2 cells **(B)** and astrocytes **(C). (D-E)** Quantified uptake of Aβ by BV-2 microglia **(D)** and astrocytes **(E)**. Units of uptake = integrated fluorescence of sample divided by the integrated fluorescence of internalized soluble Aβ42 aggregates at t_1_ for each replicate. Data points represent one of three (for astrocytes) or four (for BV-2 cells) biological replicates; error bars represent standard deviation. Statistical significance was calculated using a two-sample t-test. *P < 0.05, **P < 0.01, ***P < 0.001, ns, non-significant (P ≥ 0.05).

To isolate the role of co-aggregates, we compared the uptake of soluble aggregates from t_1_ (≥75% co-aggregates) and fibrils from t_3_ (∼0% co-aggregates) by microglia and astrocytes over a 24-hour period. Both cell types took up more soluble co-aggregates than fibrils prepared in the presence of lipidated apoE, but not non-lipidated apoE (Figure 3D-E). Uptake was also isoform-specific for soluble non-lipidated co-aggregates, which were internalized ∼2-3-fold more efficiently if they contained apoE2 rather than apoE4 (Figure S7). Co-aggregate clearance is also highly dependent on lipidation status: lipidated co-aggregates were internalized to a much greater extent than non-lipidated co-aggregates (Figure S8).

To assess how much neuroinflammation results from clearing co-aggregates, we assayed the secretion of the pro-inflammatory cytokine, TNF-α, using an enzyme-linked immunosorbent assay (ELISA) (Figure 4A). Soluble aggregates were significantly more inflammatory than fibrils if apoE was absent from the aggregation reaction or present in a non-lipidated form. However, the presence of lipidated apoE reduced the inflammation caused by soluble co-aggregates to levels similar to fibrils (Figure 4B-C). As with clearance, neuroinflammation was significantly isoform-dependent only for non-lipidated co-aggregates, which induced significantly more TNF-α secretion if they contained apoE4 rather than apoE2 (Figure S9). The lipidation status of apoE also affected neuroinflammation: co-aggregates induced higher levels of TNF-α release when they were formed in the presence of non-lipidated forms of apoE (Figure S10).

**Figure 4.**
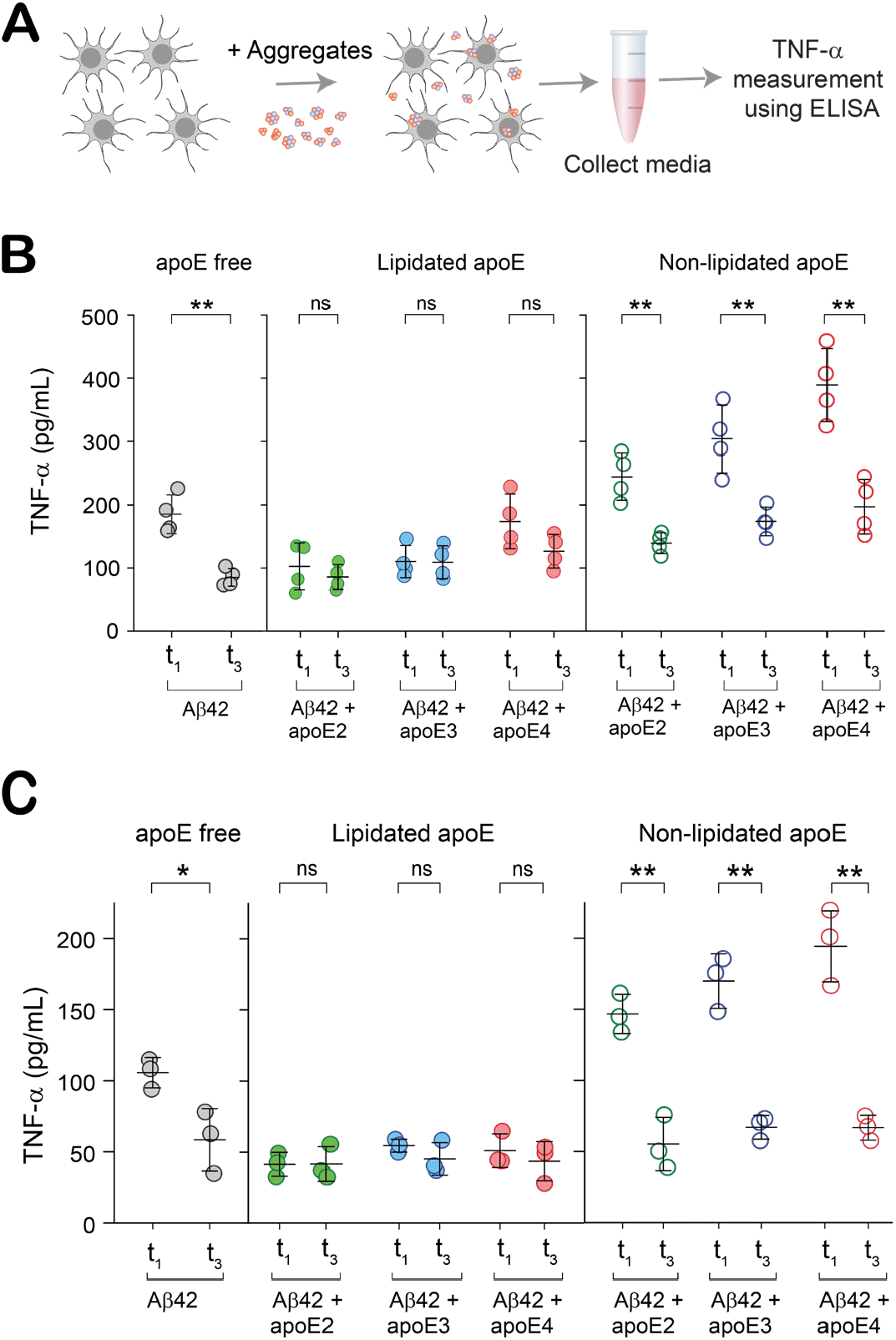
Soluble, non-lipidated apoE-Aβ co-aggregates inflame glial cells in an isoform-dependent way. **(A)** Assay for neuroinflammation of glial cells by aggregates. **(B-C)** TNF-α release by BV-2 microglia **(B)** and astrocytes **(C)** induced by soluble (t_1_) and fibrillar (t_3_) aggregates. Each data point represents one of three (for astrocytes) or four (for BV-2 cells) biological replicates; error bars represent standard deviation. Statistical significance was calculated using a two-sample t-test. *P < 0.05, **P < 0.01, ***P < 0.001, ns, non-significant (P ≥ 0.05).

These responses of microglia and astrocytes to co-aggregates begin to tease out isoform- and lipidation-dependent patterns that hint at a co-aggregate-mediated link between *APOE* genotype and AD risk. In the presence of non-lipidated apoE, soluble co-aggregates were highly inflammatory, but lipidation enhanced their uptake and suppressed inflammation. This pattern suggests a role for soluble, non-lipidated apoE-Aβ co-aggregates. Within this group, the expected isoform dependence was apparent, *i*.*e*. non-lipidated apoE4-Aβ co-aggregates were the least well cleared and most inflammatory of all Aβ species.

### Soluble, Non-lipidated apoE4-Aβ Co-aggregates are Toxic and Impair Clearance by Glial Cells

It is widely reported that aggregated Aβ is toxic, and that its toxicity depends on the size, structure, and composition of the aggregate. To check whether apoE modulates the toxicity of Aβ aggregates in an isoform or lipidation-specific way, we measured the toxicity of Aβ at different stages of aggregation. We assayed this property in two ways: i) the ability to permeabilize lipid membranes and cause Ca^2+^ influx^47^ and ii) toxicity to the SH-SY5Y human cell line, leading to the release of lactate dehydrogenase (LDH) (Figure 5A and C). We again compared soluble co-aggregates from t_1_ to fibrils from t_3_ - which have shed apoE - in order to isolate the role of co-aggregates. Soluble co-aggregates induced higher Ca^2+^ influx and LDH release than fibrils when prepared in the presence of non-lipidated apoE, but not lipidated apoE (Figure 5B and 5D). Echoing our results on uptake and neuroinflammation, the toxicity of co-aggregates was isoform-dependent (apoE4-Aβ being more toxic than apoE2-Aβ) for Aβ aggregates containing non-lipidated apoE, but not lipidated apoE (Figure S11A and S12A). Non-lipidated apoE4-Aβ co-aggregates permeabilized lipid membranes more potently (Figure S11B) and induced more LDH secretion (Figure S12B) than the corresponding lipidated forms. This lipidation dependence was absent for soluble apoE2-Aβ and apoE3-Aβ co-aggregates and fibrils prepared in the presence of all apoE isoforms.

**Figure 5.**
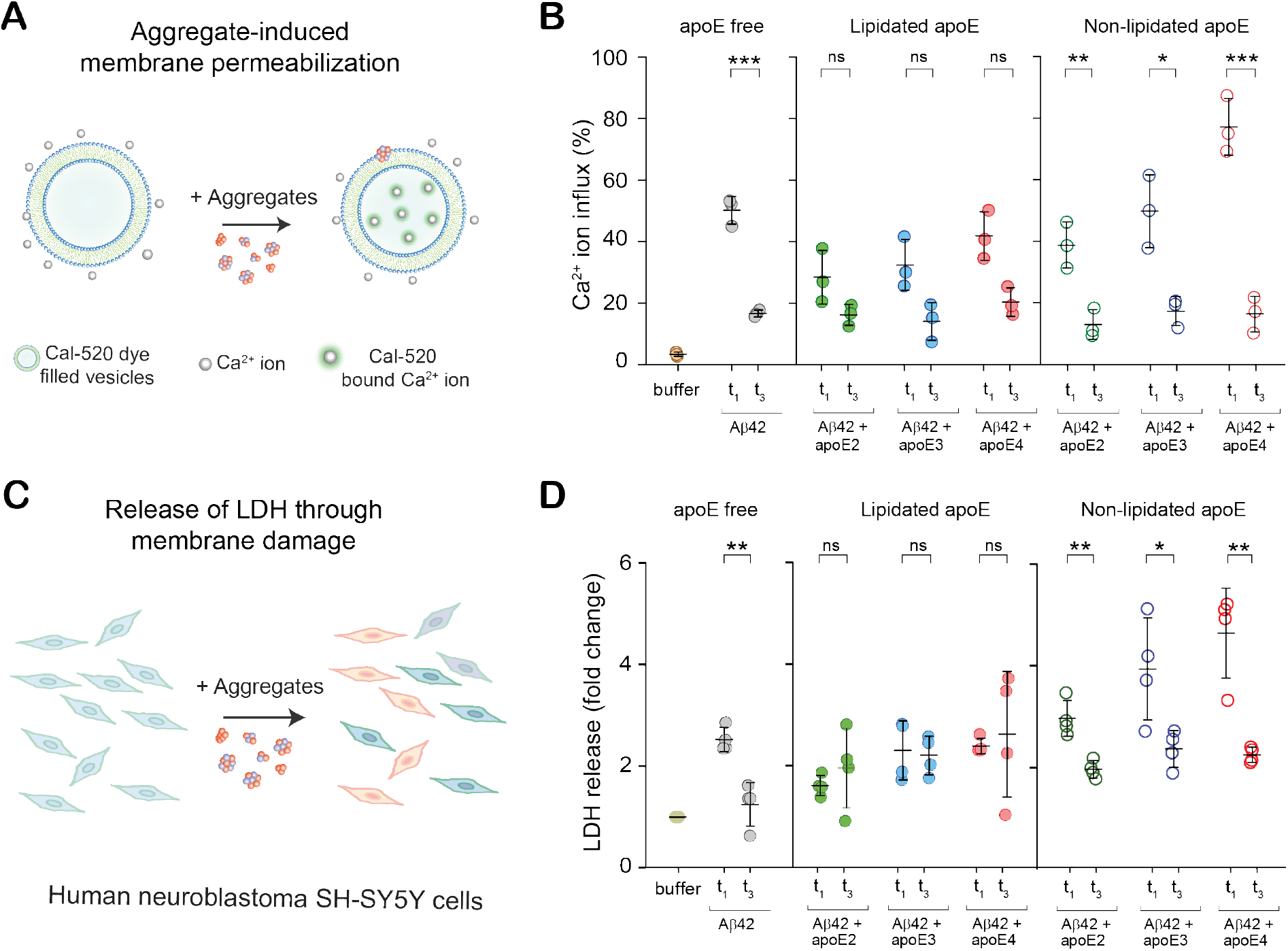
ApoE modulates the toxicity of Aβ aggregates isoform-dependently. **(A-B)** Permeabilization of lipid bilayers by soluble (t_1_) and fibrillar (t_3_) Aβ aggregates ([Aβ42] = 4 µM in monomer equivalents; [apoE] = 0 or 80 nM). Ca^2+^ influx is referenced to the influx caused by the ionophore, ionomycin. **(C-D)** Neurotoxicity of t_1_ and t_3_ aggregates to human neuroblastoma SH-SY5Y cells, assayed by lactate dehydrogenase (LDH) release ([Aβ42] = 4 µM in monomer equivalents; [apoE] = 0 or 80 nM). Data points represent one of three technical replicates **(B)** or one of three or four biological replicates **(D)**; error bars represent standard deviation. Statistical significance was calculated using a two-sample t-test. *P < 0.05, **P < 0.01, ***P < 0.001, ns, non-significant (P ≥ 0.05).

These toxicity assays painted a similar picture: soluble co-aggregates were better at permeabilizing lipid bilayers and more toxic to neuroblastoma cells than fibrils, with both lipidation- and isoform-dependent features. Lipidated co-aggregates were less toxic than non-lipidated co-aggregates, and among the latter, toxicity was isoform dependent. Non-lipidated apoE-4Aβ co-aggregates stand out as the most toxic and damaging to lipid membranes.

Having established that non-lipidated apoE-Aβ co-aggregates were especially inflammatory and toxic, we wondered whether their selective removal might rescue Aβ-induced dysfunction. To test this hypothesis, we immunoprecipitated non-lipidated apoE4-Aβ co-aggregates from a 1:1 mixture of lipidated and non-lipidated co-aggregates using HAE-4, an antibody specific for non-lipidated apoE4^15,22,25^, and an isotype control (Figure 6A-B). Cellular uptake and neuroinflammation assays on the mixture and supernatant showed that removing non-lipidated co-aggregates does indeed enhance Aβ clearance and reduces Aβ-associated inflammation in both microglia (Figure 6C-D) and astrocytes (Figure 6E-F). Immunoprecipitating non-lipidated apoE4-Aβ co-aggregates also reduced membrane damage (Figure 6G) and LDH release (Figure 6H). It is counterintuitive that removing the non-lipidated half of the aggregates increases the total uptake of Aβ (by ∼50% for astrocytes and ∼100% for microglia), even though there is ∼50% less total Aβ. However, exclusively lipidated apoE4-Aβ co-aggregates (Figure 3D-E) are taken up at well over double the levels of a 1:1 mixture of lipidated and non-lipidated co-aggregates. This implies that the inflammation caused by contact with non-lipidated co-aggregates impairs the ability of astrocytes and microglia to clear lipidated aggregates. Sparing cells from this insult by removing non-lipidated co-aggregates therefore increases the clearance of lipidated co-aggregates.

**Figure 6.**
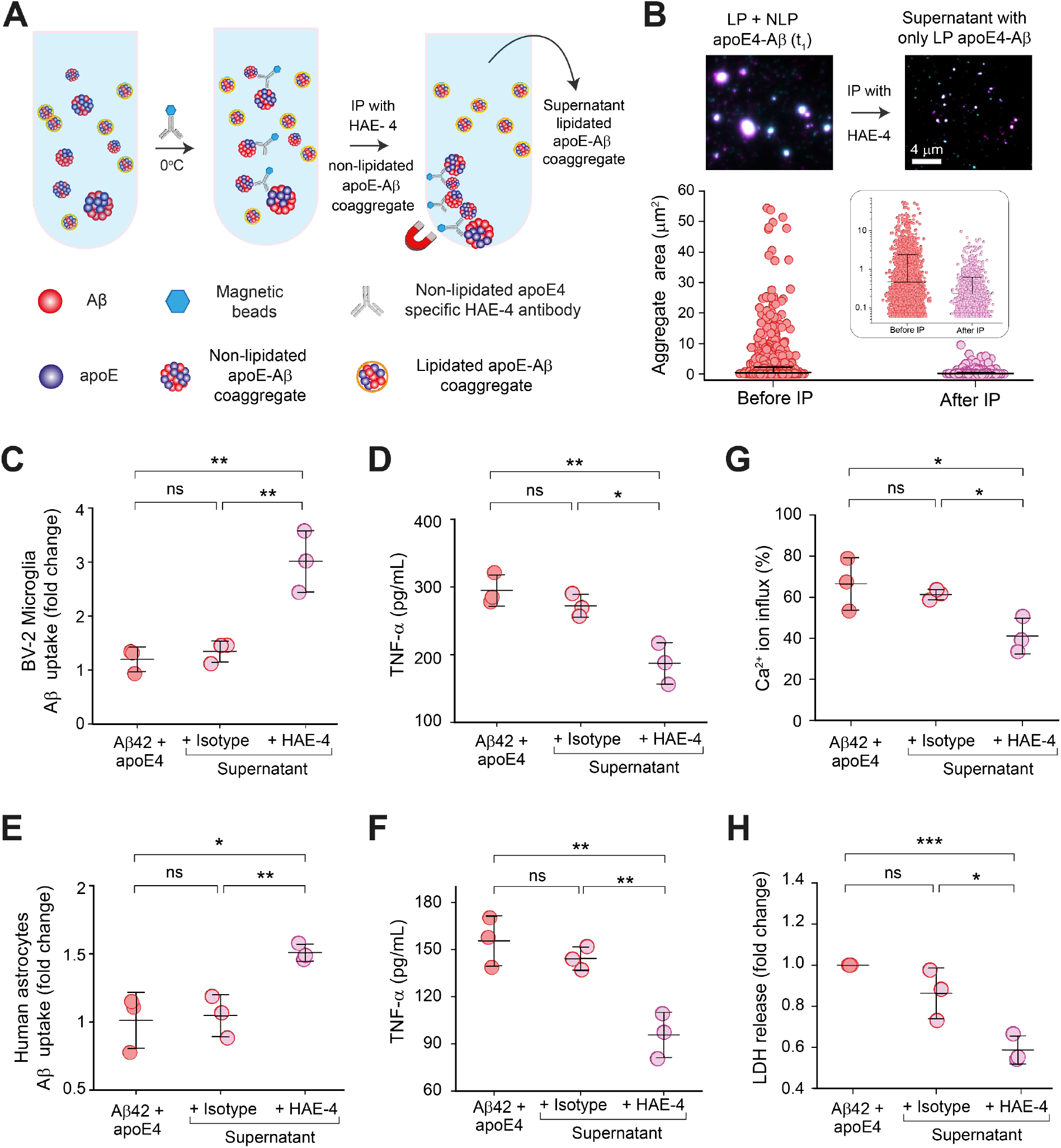
Removing non-lipidated apoE4-Aβ co-aggregates enhances Aβ clearance, reducing inflammation and cytotoxicity. **(A)** Immunoprecipitation of non-lipidated soluble co-aggregates from a mixture of non-lipidated and lipidated apoE4-Aβ co-aggregates using the HAE-4 antibody. **(B)** Two-color TIRF images and sizes of individual apoE4-Aβ co-aggregates before (number of aggregates: 10,277, average size: 0.47 ± 1.95 µm^2^) and after (number of aggregates: 4,354, average size: 0.23 ± 0.39 µm^2^) immunoprecipitation (data are plotted in log_10_ scale in inset). **(C-F)** Effect of immunoprecipitation on Aβ uptake and TNF-α release by BV-2 microglia **(C-D)** and astrocytes **(E-F). (G-H)** Effect of immunoprecipitation on neurotoxicity of apoE4-Aβ co-aggregates, measured by permeabilization of lipid membranes **(G)** and LDH release by human neuroblastoma SH-SY5Y cells **(H)**. Data points represent one of three biological replicates **(C-F** and **H)** or three technical replicates **(G)**; Units of uptake = integrated fluorescence of sample divided by the integrated fluorescence of internalized soluble Aβ42 aggregates at t_1_ for each replicate, as in Figure 3 **(C,E)**; error bars represent standard deviation. Statistical significance was calculated using One-way ANOVA with post-hoc Tukey. *P < 0.05, **P < 0.01, ***P < 0.001, ns, non-significant (P ≥ 0.05).

### Soluble apoE4-Aβ Co-aggregates Detected in AD Brains are Poorly Lipidated

Finally, to corroborate the putative role of non-lipidated apoE co-aggregates, we asked whether Aβ-aggregates containing non-lipidated apoE are over-represented in postmortem AD brain tissue. We were unable to source a specific antibody to either lipidated or non-lipidated apoE3 with sufficiently low non-specific or cross binding, so these experiments were conducted solely on samples from *APOE4/4* homozygotes. Immunoprecipitating non-lipidated apoE from these brain extracts using the HAE-4 antibody (Figure 7A) disproportionately removed the largest co-aggregates (Figure 7B). The drop in colocalization after immunoprecipitation (Figure 8C-D) implies that ∼80% of co-aggregates trapped in the neural tissue of AD patients contain poorly lipidated apoE; this is striking given that only a minority of apoE in the CNS is non-lipidated.

**Figure 7.**
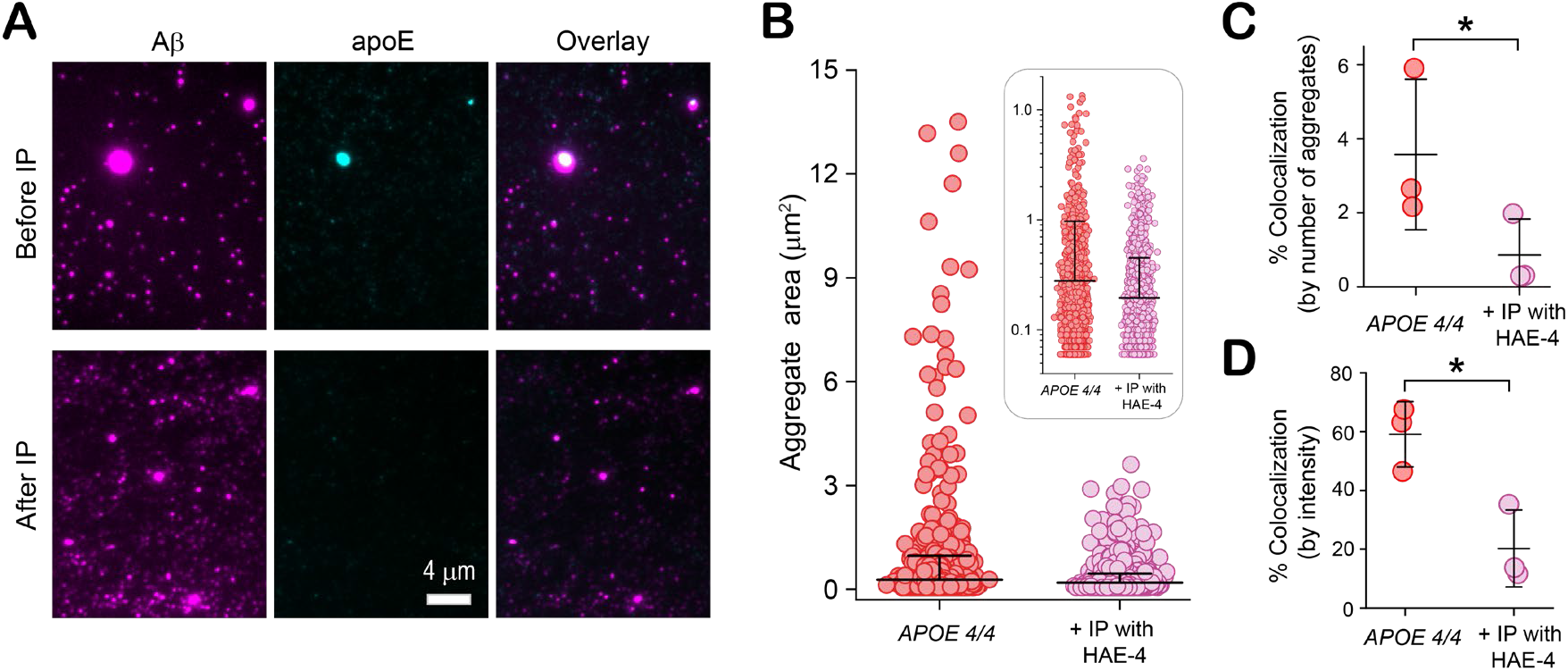
Soluble apoE4-Aβ coaggregates from the brains of *APOE4/4* AD patients are poorly lipidated. **(A)** Two-color TIRF images of soluble aggregates extracted from frontal cortices of *APOE4* homozygotes, before and after immunoprecipitation (IP) with the non-lipidated-apoE4-specific antibody, HAE-4. **(B)** Sizes of individual Aβ-containing species before (number of aggregates: 3,814, average size: 0.28 ± 0.69 µm) and after (number of aggregates: 3,576, average size: 0.19 ± 0.26 µm) immunoprecipitation (data are plotted in log_10_ scale in inset). **(C-D)** Effect of immunoprecipitation on colocalization between Aβ and apoE4 quantified by aggregate counting **(C)**, and fluorescence intensity **(D)**. Data points in panels **C** and **D** represent one of three biological replicates; error bars represent standard deviation. Statistical significance was calculated using a two-sample t-test. *P < 0.05, **P < 0.01, ***P < 0.001, ns, non-significant (P ≥ 0.05).

**Figure 8.**
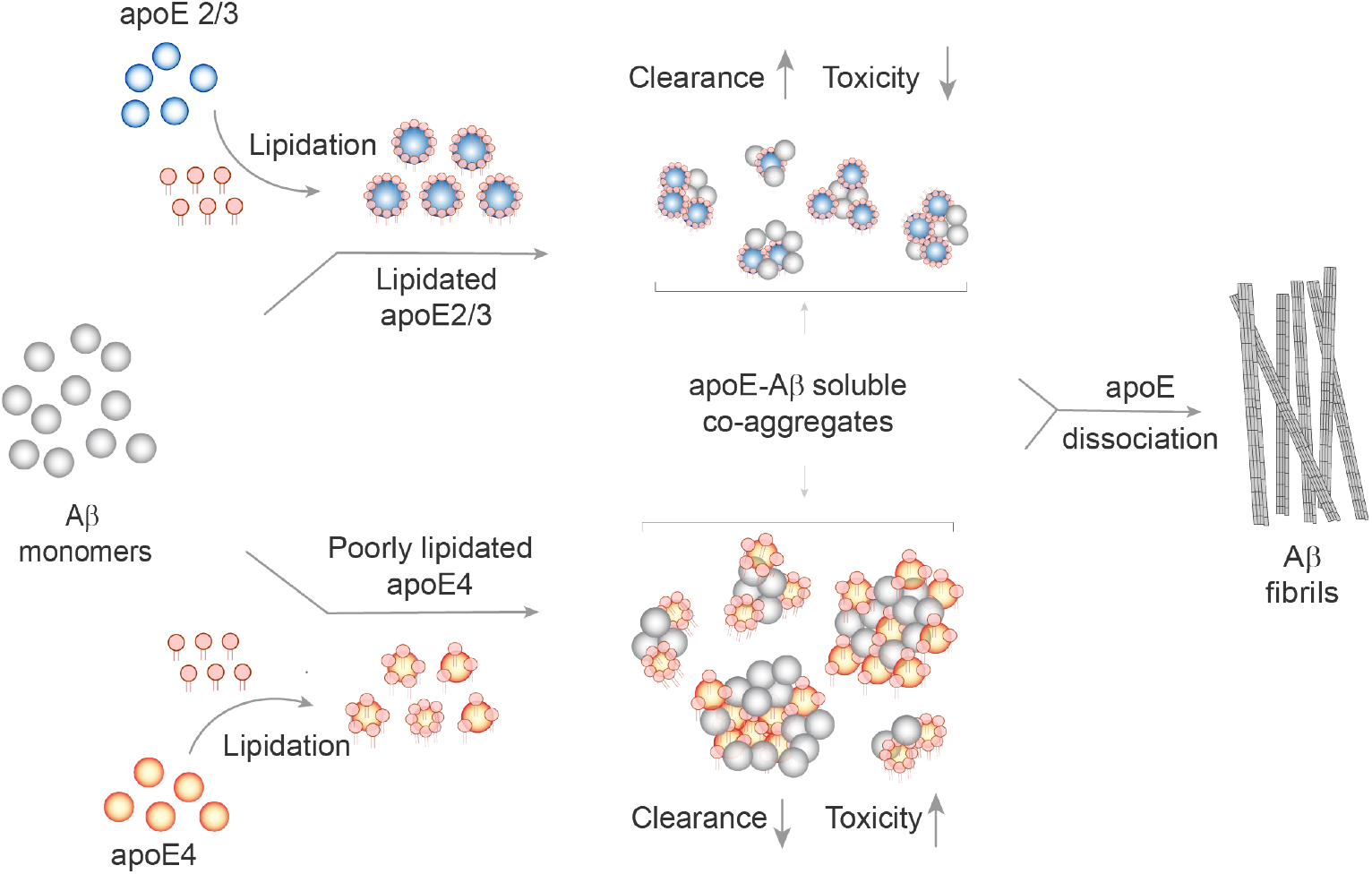
Proposed mechanism for how apoE influences Aβ aggregation and modulates Aβ clearance and toxicity in isoform- and lipidation-specific manner.

## Discussion

It is widely thought that soluble Aβ aggregates are key molecular players in AD: they trigger neuronal dysfunction, impair synaptic plasticity and their abundance correlates with severity of neurodegeneration in AD^48,49^. Attempts to reconcile this hypothesis with the link between *APOE* inheritance and late-onset AD risk have so far been unsuccessful. This study identifies a potential role for direct interactions between apoE and soluble Aβ aggregates in forging that link. Starting from the premise that apoE could affect AD risk by interacting with Aβ aggregates that are either rare or transient, we sampled a broad cross-section of the aggregation equilibrium. We found that apoE-Aβ co-aggregates are transient, but not rare: the journey to apoE-free fibrils passed through highly populated apoE-containing intermediates *in vitro*, irrespective of isoform or lipidation of apoE. In contrast, there were stark isoform- and lipidation-dependent differences in the abundances of *ex vivo* co-aggregates. Co-aggregates comprise more of the soluble aggregated Aβ detected in the frontal cortices of *APOE4/4* patients (∼50%) than *APOE3/3* patients (∼20%). About 80% of *ex vivo* co-aggregates detected from *APOE4* homozygotes contain the less-common, non-lipidated form of apoE.

Rather than deriving from aggregation kinetics or thermodynamics, the increased deposition of soluble, non-lipidated co-aggregates in *APOE4/4* brains seems to have a cellular basis. In the CNS, most Aβ aggregates are removed for degradation by glial cells *via* phagocytosis, pinocytosis, or receptor-mediated endocytosis^42,50^. Exposure to aggregates stresses microglia and astrocytes, which then release pro-inflammatory cytokines such as TNF-α, damaging nearby neurons and promoting neurodegeneration^51,52^. We found that co-aggregation with lipidated apoE enhanced Aβ uptake, while non-lipidated apoE4-Aβ co-aggregates were the slowest cleared and most inflammatory of all soluble aggregates. Our results suggest that apoE interacts with newly formed soluble aggregates in the extracellular space; if the co-aggregate contains lipidated apoE, this accelerates its clearance and avoids Aβ-induced toxicity. On the other hand, co-aggregates containing non-lipidated apoE4 are poorly cleared and inflame glial cells, which impairs the clearance of other soluble Aβ aggregates. This dynamic would lead to greater deposition of Aβ over time in individuals with *APOE4* genotypes. The fact that apoE4 is less lipidated *in vivo*^53,54^ than apoE2 or apoE3 may exacerbate this vicious circle.

This apoE-mediated mechanism for Aβ deposition, combined with co-aggregate toxicity may influence disease progression in AD. Non-lipidated apoE-Aβ co-aggregates are more likely to be deposited than lipidated co-aggregates, but are also more toxic. This may explain why deleting the ATP-binding cassette transporter 1 (ABCA1) - which regulates apoE lipidation in the CNS^16^ – increases the deposition of Aβ plaques^55,56^, while overexpressing or upregulating ABCA1 relieves cognitive impairment caused by Aβ and reverses memory deficits in transgenic animals^57^. The effect of lipidation, alongside the isoform-dependence of co-aggregate toxicity connect the *APOE4* genotype with increased amyloid load, as well as faster AD onset and progression, *via* a possible culprit: soluble, non-lipidated apoE4-Aβ co-aggregates. Importantly, mature Aβ fibrils do not contain apoE and are functionally indistinguishable, whichever form of apoE was present during aggregation. This finding strengthens our hypothesis that apoE contributes to AD risk by modulating the behaviour of soluble Aβ aggregates.

Our work suggests further avenues for investigating and treating late-onset AD. We show that removing non-lipidated apoE4 co-aggregates reduces inflammation, enhances Aβ uptake by glial cells, and protects from Aβ-induced membrane disruption and subsequent Ca^2+^ dysregulation. This may form the molecular basis of immunotherapy with the HAE-4 monoclonal antibody raised against non-lipidated human apoE^15,22^. We found opposing trends in the isoform-dependence of internalization and membrane permeabilization of co-aggregates, suggesting that uptake may be mainly receptor-mediated. Because the receptor binding capacity of apoE depends on its lipidation^58,59^, the fact that lipidation of co-aggregates enhances their uptake supports this idea. ApoE’s influence on endocytosis is well studied, with roles for other AD risk factors such as LDL^60^, TREM2^61^ and *PICALM*^62^, while LRP1 and VLDLR are putative receptors for the transport of apoE-Aβ complexes^27^. Validating specific receptor(s) which play a significant role in the internalization of large soluble co-aggregates could present opportunities to precisely target apoE in AD. Modulating the formation or uptake of non-lipidated, soluble co-aggregates in an isoform-specific way could eliminate the AD-specific role of apoE without impeding its essential functions in lipid transport. Enhancing apoE lipidation could also provide a new intervention strategy for late-onset Alzheimer’s disease, specifically for *APOE4* carriers, because apoE4 is less lipidated than apoE2 or apoE3^5,54^.

## Limitations of study

This work advances a mechanism for how apoE isoform and lipidation influence neurodegeneration in Alzheimer’s disease *via* transient, soluble apoE-Aβ co-aggregates. However, several questions remain unanswered. The prevalence of soluble apoE-Aβ coaggregates in the post-mortem tissue of AD patients provides strong circumstantial support for our hypothesis, but does not rule out contributions from other mechanisms such as neuronal susceptibility to injury, insulin signalling, or neuroinflammation *via* overactivation of the nuclear factor kappa B. Developing a means to isolate and functionally study these co-aggregates over the course of AD would provide an opportunity to test our hypothesis further. We also do not yet understand why soluble apoE-Aβ co-aggregates differ in size and their ability to damage lipid membranes. We hypothesize that this is because co-aggregates differ in structure and/or stoichiometry, though we do not have definitive proof. New techniques will be required to study the composition and high-resolution structure of soluble aggregates, and hence reveal structure-function relationships at the single-particle level. Finally, we have not yet identified the molecular basis for the isoform-dependence of apoE-Aβ coaggregate uptake by microglia and astrocytes. Future work will need to determine what role(s) apoE and Aβ binding receptors play in the differential clearance of co-aggregates.

## Author Contributions

Z.X, E.E.P., R.T.R. and S.D. did the SiMPull experiments. R.T.R. and E.E.P. modified antibodies and coated coverslips for SiMPull. Z.X., H.E.W and S.D. aggregated proteins. The cellular uptake assay was carried out by M.M., E.E.P and H.E.W. A.H. helped with the uptake data analysis. H.D. and S.D. prepared soaked-brain extracts. Z.X., E.E.P., and S.D. did the membrane permeabilization and LDH assays. H.J., H.Z., Y.L.J.L., Y.Z., and J.D. helped with antibody modifications, slide preparation or microscopy. Data analysis and statistics were developed, carried out by Z.X., and interpreted by Z.X. and S.D. L.F., S.W., S.M.B., R.T.R., A.H., H.M., P.T., D.C.C., D.M.H., D.K., and S.D. supervised the project. The initial draft of the paper was written by R.T.R., D.K., E.E.P. and S.D.; all other authors provided feedback and contributed in editing the manuscript into its final form. D.K. and S.D. designed and conceived the study. All authors read and approved the manuscript.

## Acknowledgments

The tissue for this study was provided by the Newcastle Brain Tissue Resource, which is funded in part by a grant from the UK Medical Research Council and in part by Brains for Dementia Research, a joint venture between Alzheimer’s Society and Alzheimer’s Research UK. We also thank the Professor Maria Spillantini laboratory for providing the facilities and expertise for processing of the brain tissue used in these experiments.

## Resource Availability

### Data availability

All data reported in this paper will be shared by the lead contact upon request.

### Lead contact

Further information and requests for reagents may be directed to and will be fulfilled by the lead contact Suman De.

## References

1. Liu,C. C., Kanekiyo, T., Xu, H. & Bu, G. Apolipoprotein E and Alzheimer disease: risk, mechanisms and therapy. Nat. Rev. Neurol. 9, 106–18 (2013).

2. Conejero-Goldberg, C. et al. APOE2 enhances neuroprotection against Alzheimer’s disease through multiple molecular mechanisms. Mol. Psychiatry 19, 1243–1250 (2014).

3. Corder, E. H. et al. Gene dose of apolipoprotein E type 4 allele and the risk of Alzheimer’s disease in late onset families. Science (80-.). 261, 921–923 (1993).

4. Verghese, P. B., Castellano, J. M. & Holtzman, D. M. Apolipoprotein E in Alzheimer’s disease and other neurological disorders. Lancet Neurol. 10, 241–252 (2011).

5. Kanekiyo, T., Xu, H. & Bu, G. ApoE and Aβ; in Alzheimer’s Disease: Accidental Encounters or Partners? Neuron 81, 740–754 (2014).

6. Hardy, J. & Selkoe, D. J. The Amyloid Hypothesis of Alzheimer’s Disease: Progress and Problems on the Road to Therapeutics. Science (80-.). 297, 353–356 (2002).

7. Benilova, I., Karran, E. & De Strooper, B. The toxic Aβ oligomer and Alzheimer’s disease: an emperor in need of clothes. Nat. Neurosci. 15, 349–57 (2012).

8. Li, S. et al. Soluble oligomers of amyloid β protein facilitate hippocampal long-term depression by disrupting neuronal glutamate uptake. Neuron 62, (2009).

9. De Strooper, B. et al. Deficiency of presenilin-1 inhibits the normal cleavage of amyloid precursor protein. Nature 391, 387–390 (1998).

10. Masters, C. L. et al. Amyloid plaque core protein in Alzheimer disease and Down syndrome. Proc. Natl. Acad. Sci. 82, 4245–4249 (1985).

11. Choi, S. H. et al. A three-dimensional human neural cell culture model of Alzheimer’s disease. Nature 515, 274–278 (2014).

12. Gonneaud, J. et al. Relative effect of APO ε4 on neuroimaging biomarker changes across the lifespan. Neurology 87, 1696–1703 (2016).

13. Fleisher, A. S. et al. Apolipoprotein E ε4 and age effects on florbetapir positron emission tomography in healthy aging and Alzheimer disease. Neurobiol. Aging 34, 1–12 (2013).

14. Liu, C.-C. et al. ApoE4 Accelerates Early Seeding of Amyloid Pathology. Neuron 96, 1024-1032.e3 (2017).

15. Liao, F. et al. Targeting of nonlipidated, aggregated apoE with antibodies inhibits amyloid accumulation. J. Clin. Invest. 128, 2144–2155 (2018).

16. Wahrle, S. E. et al. ABCA1 Is Required for Normal Central Nervous System ApoE Levels and for Lipidation of Astrocyte-secreted apoE*. J. Biol. Chem. 279, 40987–40993 (2004).

17. Mouchard, A. et al. ApoE-fragment/Aβ heteromers in the brain of patients with Alzheimer’s disease. Sci. Rep. 9, 3989 (2019).

18. Mormino, E. C. et al. Amyloid and APOE ε4 interact to influence short-term decline in preclinical Alzheimer disease. Neurology 82, 1760–1767 (2014).

19. Lim, Y. Y. et al. APOE and BDNF polymorphisms moderate amyloid β-related cognitive decline in preclinical Alzheimer’s disease. Mol. Psychiatry 20, 1322–1328 (2015).

20. Kantarci, K. et al. APOE modifies the association between Aβ load and cognition in cognitively normal older adults. Neurology 78, 232–240 (2012).

21. Castellano, J. M. et al. Human apoE Isoforms Differentially Regulate Brain Amyloid-β Peptide Clearance. Sci. Transl. Med. 3, 89ra57 (2011).

22. Xiong, M. et al. APOE immunotherapy reduces cerebral amyloid angiopathy and amyloid plaques while improving cerebrovascular function. Sci. Transl. Med. 13, eabd7522 (2021).

23. Näslund, J. et al. Characterization of stable complexes involving apolipoprotein E and the amyloid β peptide in Alzheimer’s disease brain. Neuron 15, 219–228 (1995).

24. Huynh, T.-P. V, Davis, A. A., Ulrich, J. D. & Holtzman, D. M. Apolipoprotein E and Alzheimer’s disease: the influence of apolipoprotein E on amyloid-β and other amyloidogenic proteins: Thematic Review Series: ApoE and Lipid Homeostasis in Alzheimer’s Disease. J. Lipid Res. 58, 824–836 (2017).

25. Gratuze, M. et al. APOE Antibody Inhibits Aβ-Associated Tau Seeding and Spreading in a Mouse Model. Ann. Neurol. 91, 847–852 (2022).

26. Verghese, P. B. et al. ApoE influences amyloid-β (Aβ) clearance despite minimal apoE/Aβ association in physiological conditions. Proc. Natl. Acad. Sci. 110, 1807–1816 (2013).

27. Deane, R. et al. apoE isoform–specific disruption of amyloid β peptide clearance from mouse brain. J. Clin. Invest. 118, 4002–4013 (2008).

28. Ono, K., Condron, M. M. & Teplow, D. B. Structure–neurotoxicity relationships of amyloid β-protein oligomers. Proc. Natl. Acad. Sci. 106, 14745–14750 (2009).

29. De, S. et al. Different soluble aggregates of Aβ42 can give rise to cellular toxicity through different mechanisms. Nat. Commun. 10, 1541 (2019).

30. Xian, X. et al. Reversal of ApoE4-induced recycling block as a novel prevention approach for Alzheimer’s disease. Elife 7, e40048 (2018).

31. Minta, K. et al. Quantification of total apolipoprotein E and its isoforms in cerebrospinal fluid from patients with neurodegenerative diseases. Alzheimers. Res. Ther. 12, 19 (2020).

32. Galvagnion, C. et al. Lipid vesicles trigger α-synuclein aggregation by stimulating primary nucleation. Nat. Chem. Biol. 11, 229–234 (2015).

33. Tahirbegi, B. et al. A Novel Aβ40 Assembly at Physiological Concentration. Sci. Rep. 10, 9477 (2020).

34. Lindberg, D. J., Wesén, E., Björkeroth, J., Rocha, S. & Esbjörner, E. K. Lipid membranes catalyse the fibril formation of the amyloid-β (1–42) peptide through lipid-fibril interactions that reinforce secondary pathways. Biochim. Biophys. Acta -Biomembr. 1859, 1921–1929 (2017).

35. Jain, A. et al. Probing cellular protein complexes using single-molecule pull-down. Nature 473, 484–488 (2011).

36. Cerf, E., Gustot, A., Goormaghtigh, E., Ruysschaert, J.-M. & Raussens, V. High ability of apolipoprotein E4 to stabilize amyloid-β peptide oligomers, the pathological entities responsible for Alzheimer’s disease. FASEB J. 25, 1585–1595 (2011).

37. Küffner, A. M. et al. Sequestration within biomolecular condensates inhibits Aβ-42 amyloid formation. Chem. Sci. 12, 4373–4382 (2021).

38. Hong, W. et al. Diffusible, highly bioactive oligomers represent a critical minority of soluble Aβ in Alzheimer’s disease brain. Acta Neuropathol. 136, 19–40 (2018).

39. Sideris, D. I. et al. Soluble amyloid beta-containing aggregates are present throughout the brain at early stages of Alzheimer’s disease. Brain Commun. 3, fcab147 (2021).

40. Koffie, R. M. et al. Apolipoprotein E4 effects in Alzheimer’s disease are mediated by synaptotoxic oligomeric amyloid-β. Brain 135, 2155–2168 (2012).

41. Hashimoto, T. et al. Apolipoprotein E, Especially Apolipoprotein E4, Increases the Oligomerization of Amyloid β Peptide. J. Neurosci. 32, 15181 LP – 15192 (2012).

42. Paresce, D. M., Ghosh,R. N. & Maxfield, F. R. Microglial Cells Internalize Aggregates of the Alzheimer’s Disease Amyloid β-Protein;-Protein Via a Scavenger Receptor. Neuron 17, 553–565 (1996).

43. Liu, C.-C. et al. Astrocytic LRP1 Mediates Brain Aβ Clearance and Impacts Amyloid Deposition. J. Neurosci. 37, 4023 LP – 4031 (2017).

44. Hickman, S. E., Allison, E. K. & El Khoury, J. Microglial Dysfunction and Defective β-Amyloid Clearance Pathways in Aging Alzheimer’s Disease Mice. J. Neurosci. 28, 8354 LP – 8360 (2008).

45. Garwood, C. J., Pooler, A. M., Atherton, J., Hanger, D. P. & Noble, W. Astrocytes are important mediators of Aβ-induced neurotoxicity and tau phosphorylation in primary culture. Cell Death Dis. 2, e167–e167 (2011).

46. Craft, J. M., Watterson, D. M. & Van Eldik, L. J. Human amyloid β-induced neuroinflammation is an early event in neurodegeneration. Glia 53, 484–490 (2006).

47. Flagmeier, P. et al. Direct measurement of lipid membrane disruption connects kinetics and toxicity of Aβ42 aggregation. Nat. Struct. Mol. Biol. 10, 886–891 (2020).

48. Hsiao, K. et al. Correlative Memory Deficits, Aβ Elevation, and Amyloid Plaques in Transgenic Mice. Science (80-.). 274, 99–103 (1996).

49. McLean, C. A. et al. Soluble pool of Aβ amyloid as a determinant of severity of neurodegeneration in Alzheimer’s disease. Ann Neurol 46, 860–866 (1999).

50. Mandrekar, S. et al. Microglia Mediate the Clearance of Soluble Aβ through Fluid Phase Macropinocytosis. J. Neurosci. 29, 4252 LP – 4262 (2009).

51. Wang, W.-Y., Tan, M.-S., Yu, J.-T. & Tan, L. Role of pro-inflammatory cytokines released from microglia in Alzheimer’s disease. Ann. Transl. Med. Vol 3, No 10 (June 2015) Ann. Transl. Med. (2015).

52. Griffin, W. S. T. et al. Glial-Neuronal Interactions in Alzheimer’s Disease: The Potential Role of a ‘Cytokine Cycle’ in Disease Progression. Brain Pathol. 8, 65–72 (1998).

53. Hanson, A. J. et al. Effect of Apolipoprotein E Genotype and Diet on Apolipoprotein E Lipidation and Amyloid Peptides: Randomized Clinical TrialApolipoprotein E Lipidation and Amyloid Peptides. JAMA Neurol. 70, 972–980 (2013).

54. Hu, J. et al. Opposing effects of viral mediated brain expression of apolipoprotein E2 (apoE2) and apoE4 on apoE lipidation and Aβ metabolism in apoE4-targeted replacement mice. Mol. Neurodegener. 10, 6 (2015).

55. Wahrle, S. E. et al. Deletion of Abca1 Increases Aβ Deposition in the PDAPP Transgenic Mouse Model of Alzheimer Disease. J. Biol. Chem. 280, 43236–43242 (2005).

56. Koldamova, R., Staufenbiel, M. & Lefterov, I. Lack of ABCA1 Considerably Decreases Brain ApoE Level and Increases Amyloid Deposition in APP23 Mice. J. Biol. Chem. 280, 43224–43235 (2005).

57. Wahrle, S. E. et al. Overexpression of ABCA1 reduces amyloid deposition in the PDAPP mouse model of Alzheimer disease. J. Clin. Invest. 118, 671–682 (2008).

58. Frieden, C., Wang, H. & Ho, C. M. W. A mechanism for lipid binding to apoE and the role of intrinsically disordered regions coupled to domain–domain interactions. Proc. Natl. Acad. Sci. 114, 6292–6297 (2017).

59. Chen, Y., Strickland, M. R., Soranno, A. & Holtzman, D. M. Apolipoprotein E: Structural Insights and Links to Alzheimer Disease Pathogenesis. Neuron 109, 205–221 (2021).

60. Holtzman, D. M., Herz, J. & Bu, G. Apolipoprotein E and Apolipoprotein E Receptors: Normal Biology and Roles in Alzheimer Disease. Cold Spring Harb. Perspect. Med. 2, (2012).

61. Yeh, F. L., Wang, Y., Tom, I., Gonzalez, L. C. & Sheng, M. TREM2 Binds to Apolipoproteins, Including APOE and CLU/APOJ, and Thereby Facilitates Uptake of Amyloid-Beta by Microglia. Neuron 91, 328–340 (2016).

62. Narayan, P. et al. PICALM Rescues Endocytic Defects Caused by the Alzheimer’s Disease Risk Factor APOE4. Cell Rep. 33, 108224 (2020).

