## Supplemental Figures and Methods for "Co-aggregation with Apolipoprotein E modulates the function of Amyloid-β in Alzheimer’s disease"

**Methods and supplementary information**

### Materials and methods

**Aggregation of recombinant Aβ42:** Solutions of monomeric recombinant Aβ42 peptide (Stratech, Cat. No. A-1167-2) were diluted in phosphate-buffered saline (PBS) buffer to a concentration of 4 μM and aggregated in a 96-well half-area plate (Corning, Cat. No. 3881) at 37°C, under quiescent conditions in the absence and presence of non-lipidated or lipidated apolipoprotein E (apoE) isoforms E2, E3 or E4 (Peprotech, Cat. No. 350-12, 350-02 or 350-04) at concentrations of 80 nM (lipidated apoE2, apoE3 and apoE4, and non-lipidated apoE2 and apoE3) or 20, 40 and 80 nM (non-lipidated apoE4). The aggregation process was monitored using 20 μM Thioflavin T dye (Sigma, Cat. No. T3516) using a plate reader (Clariostar Plus, BMG Biotech).

To analyze the species formed during Aβ42 aggregation and co-aggregation with apoE, samples of the aggregation mixture (aggregated without ThT) were taken at the end of the lag phase ($t_1$), ~15 minutes' incubation (30 min and 10 min for co-aggregation with non-lipidated and lipidated apoE respectively), middle of the growth phase ($t_2$) ~30 minutes' incubation (45 min and 20 min for co-aggregation with non-lipidated and lipidated apoE respectively) and at the plateau phase ($t_3$) ~90 minutes' incubation (120 min for co-aggregation with both non-lipidated and lipidated apoE).

**Lipidation of recombinant apoE:** ApoE was lipidated using a previously published protocol[1]. All the isoforms of apoE were lipidated using a cholate dialysis method using 1-palmitoyl-2-oleoyl-glycerol-3-phosphocholine 16:0-18:1 PC (POPC) (Avanti Lipids, 850457C) and Cholesterol (Avanti Lipids, 700100P) in a molar ratio 1:90:5 (apoE: POCP: cholesterol). These ratios of apoE and lipids were selected to mimic the physiological lipid composition of HDL-like apoE particles. Lipids can be directly incorporated into recombinant apoE, but the sodium cholate dialysis method has been shown to produce lipidated apoE homogeneously in a reproducible manner.

First, recombinant apoE was suspended in 600 μL of apoE buffer [20 mM phosphate buffer (Sigma) containing 50 mM NaCl (Sigma, Cat. No. S7653), 1 mM dithiothreitol (DTT, Sigma, Cat. No. 10708984001), and 1 mM ethylenediaminetetraacetic acid (EDTA, Sigma, Cat. No. E9884) at pH 7.4]. In parallel, stock solutions of 10 mg/mL 16:0-18:1 PC (Avanti Lipids, Cat. No. 850457C), and 10 mg/mL cholesterol (Avanti Lipids, Cat. No. 700100P) in chloroform were made. Then 16:0-18:1 PC (18.3 μL) and cholesterol (2 μL) were mixed in a glass vial at a molar ratio of 90:5 and the chloroform was then removed under vacuum in a desiccator overnight. Then 40 μL of Dulbecco's phosphate-buffered saline buffer (DPBS, Thermo Fisher, Cat. No. 141901440) with 0.5 mM DTT (Sigma) was added to each vial to resuspend the lipid at a total lipid concentration of 5 μg/μL, which was then vortexed three times for 10 min at 2 minute intervals. Then the solution left to stand for 30 minutes at room temperature. Then, sodium cholate (50 mg·mL$^{-1}$, Sigma) was slowly added to the lipid solutions with vortexing until the solution was free of turbidity. The apoE solution was then added and the mixture vortexed three times for 10 minutes, at 5 minute intervals and left to stand for 1 hour. Then the entire solution was dialyzed using Slide-A-Lyzer Dialysis Cassettes with a 10 kDa -cut off (Thermo Fisher Cat 66380) with apoE buffer for 72 h, (with a buffer change every 24 h) at 4°C. After this, the apoE solutions were purified by gel filtration chromatography (Superose 6 Increase 3.2/300) at 4°C, and the fractions containing lipidated apoE were combined and concentrated using an Amicon Ultra 15 mL 10 kDa cut-off centrifugal filter unit (Merck Millipore, Cat. No. UFC901024) at 3000 g for 20 min at 4°C. The concentration of lipidated apoE was determined by absorbance measurements at 280 nm using an extinction coefficient of 44,460 M$^{-1}$·cm$^{-1}$. Finally, the samples of lipidated apoE were stored at 4°C. Lipidation of all the apolipoproteins were performed in parallel using the same lipid–cholesterol suspension.

**Postmortem brain tissue:** Flash-frozen brain tissues from six Alzheimer's disease patients (three homozygous *APOE4* and three homozygous *APOE3* AD patients) were obtained from the Newcastle brain tissue resource (NTBR) under a Material Transfer Agreement. Tissue was processed at the NTBR, where regions of interest were removed from the frontal cortex (Brodmann area 9) and immediately frozen at -80°C. Data relating to supplied tissue were released on an anonymized basis. Ethical approval was granted through the Newcastle and North Tyneside Research Ethics Committee.

**Supplementary Table 1.** Properties of postmortem brain tissue samples.

| Sex | Age | Postmortem time before freezing (hours) | Braak stage | *APOE* genotype |
|---|---|---|---|---|
| F | 70 | 24 | 6 | 4/4 |
| M | 90 | 69 | 6 | 4/4 |
| F | 80 | 10 | 6 | 4/4 |
| F | 58 | 70 | 6 | 3/3 |
| F | 84 | 26 | 6 | 3/3 |
| F | 88 | 57 | 6 | 3/3 |

**Extraction of soluble protein aggregates from postmortem tissue:** Soluble Aβ-containing aggregates were extracted from human tissue following a recently published protocol[2] with minor changes. In summary, the supplied tissues were chopped into 300 mg pieces using a razor blade and incubated with 1 mL of artificial cerebrospinal fluid (aCSF) buffer (120 mM NaCl, 2.5 mM KCl, 1.5 mM $NaH_2PO_4$, 26 mM $NaHCO_3$, 1.3 mM $MgCl_2$, pH 7.4) at 4°C for 30 min under gentle agitation. The mixture was then centrifuged for 10 min at 2,000 g at 4°C. The upper ~80% of the supernatant was further centrifuged for 110 min at 14,000 g at 4°C. The upper ~80% of the supernatant was collected and dialyzed using a 2 kDa molecular-weight-cut-off (MWCO) Slide-A-Lyzer (Thermo Fisher) for 72 hours at 4°C, against a 50-fold excess of aCSF buffer, which was changed every 12 hours. These samples were aliquoted into small volumes and stored at -80°C, and used for further experiments with no further freeze-thaw cycles.

**Materials for SiMPull assays:** Biotinylated 6E10 (Biolegend, Cat. No. 803007, Lot No. B230416), Alexa-Fluor-647-labeled 6E10 (Biolegend, Cat. No. 803021, Lot No. B304121), Alexa-Fluor-488-labeled anti-apoE antibody F-9 (Santa Cruz Biology, Cat. No. sc-390925 AF488, Lot No. B1717), Alexa-Fluor-647-labeled Mouse IgG1 (isotype control, MGL, Cat. No. M075-A64, Lot. No. 002), Alexa-Fluor-488-labeled Mouse IgG (isotype control, Fisher Scientific, Cat. No. 65-0865-14) antibodies were used without further modification or purification and stored at 4°C until needed. Anti-apoE antibody EPR19392 (Abcam, Cat. No. ab227993, Lot No. GR3268327-5,130 µg, at 1.3 mg/mL in PBS) was mixed with twenty molar equivalents of Alexa Fluor 488 TFP ester (17.2 nmol, 8 mM in dimethyl sulfoxide) and 10 µL of 1 M $NaHCO_3$, and incubated for 2 h. The excess dye was removed using a Zeba Spin Desalting Column (7 kDa MWCO), then an Amicon spin filter (50 kDa MWCO), and the concentration of antibody and degree of labeling (~1.5 dyes per antibody) were measured using A495 and A280 (Nanodrop, Thermo Fisher). The labeled antibody was diluted to 0.5 mg/mL in 0.5 x PBS containing 0.01% sodium azide and 50% glycerol and stored at -20°C until needed. NeutrAvidin (Thermo Fisher, Cat. No. 31000) and bovine serum albumin (BSA, Thermo Fisher, Cat. No. 10829410) were used without further purification and stored at 4°C and -20°C, respectively. Glass coverslips (26x76 mm, thickness #1.5, VWR, Cat. No. MENZBC026076AC40) were covalently PEGylated precisely as we reported previously[3] and stored in a desiccator at -20°C until needed.

**SiMPull protocol:** Assay wells were first coated with 10 µL of 0.2 mg/mL NeutrAvidin in 1× PBS containing 0.05% Tween-20 for 5 min, washed twice with 10 µL of 1× PBS containing 0.05% Tween-20, and once with 10 µL of 1× PBS containing 1% Tween-20. Ten microliters of biotinylated 6E10 (10 nM in 1× PBS containing 0.1 mg/mL BSA) were added to each well for 15 min. The wells were then washed twice with 10 µL of 1× PBS containing 0.05% Tween-20, and once with 10 µL of 1× PBS containing 1% Tween-20. The (co-)aggregates taken at different time points (diluted to 1 µM monomer equivalents in PBS) or soaked-brain extracts (neat) were then added and incubated for 1 hour. The wells were then washed twice with 10 µL of 1× PBS containing 0.05% Tween-20, and once with 10 µL of 1× PBS containing 1% Tween-20. The mixture of imaging antibodies (1 nM Alexa-Fluor-488-labeled EPR19392 and 500 pM Alexa-Fluor-647-labeled 6E10 in 1× PBS containing 0.1 mg/mL BSA) were then added and incubated for 30 min. The wells were then washed twice with 10 µL of 1× PBS containing 0.05% Tween-20, and once with 10 µL of 1× PBS containing 1% Tween-20. Finally, 3 µL of 1× PBS were added to each well and samples were sealed using a second plasma-cleaned coverslip.

**Imaging setup for SiMPull:** Samples were imaged using a custom-built total internal reflection fluorescence (TIRF) microscope based on a Nikon Ti2 microscope fitted with a Perfect Focus unit. Laser beams (488 nm and 635 nm, Oxxius) were circularly polarized using a quarter-wave plate (WPQ05M-405, Thorlabs), then focused onto the back focal plane of a 100x Plan Apo TIRF, NA 1.49 oil-immersion objective lens (Nikon). Fluorescence emission was collected by the same objective and separated from the excitation beam using a dichroic beamsplitter (Di01-R405/488/561/635, Laser 2000). The emitted light was passed through a 488 nm long-pass filter (BLP01-488R, Laser 2000) and bandpass filter (FF01-520/44-25, Laser 2000) for Alexa-Fluor-488 emission, and a 635 nm long-pass filter (BLP01-635R-25, Laser 2000) for Alexa-Fluor-647 emission and imaged on an EMCCD camera (Photometrics Evolve). The open-source software Micro-Manager 1.4 was used to automate image acquisition[4]. Images were acquired in an unbiased way at laser powers of ~10 W/cm$^2$ and averaged over 50 frames with an exposure time of 50 ms each.

**Analysis of SiMPull data:** The averaged images acquired with excitation at 488 nm (apolipoprotein E detection channel) and 635 nm (amyloid-beta detection channel) were passed to the Fiji plugin ComDet3[5] with a self-written python automation code. Particles are differentiated from noise by selecting 20 times intensity significance (20x SD) compared to the background. For spots in two different channels to be considered colocalized, the displacement between their centers of mass (determined by Gaussian fitting) was required to be ≤3 pixels. Colocalization by aggregate number was defined as the ratio between the number of colocalized spots and the total number of spots in the amyloid-beta channel. Colocalization by intensity was defined as the ratio between the sum of the integrated intensities of colocalized spots and all spots in the amyloid-beta channel. Random colocalization was estimated by performing the colocalization algorithm with one of the images (50 µm x 50 µm) rotated by 90 degrees, which was less than 1% for all data sets.

**BV-2 Microglia Culture and immunohistochemistry:** BV-2 cells were maintained in Dulbecco's modified Eagles medium (DMEM) Glutamax (Thermo Fisher, Cat. No. 10569010), supplemented with 10% fetal bovine serum (FBS) (Thermo Fisher, Cat. No. 26140087), 100 U/mL penicillin and 100 µg/mL streptomycin (P/S) (Thermo Fisher, Cat. No. 15140148) and incubated at 37°C in 5% $CO_2$ and 95% air. Cells were grown to ~80% confluency and sub-cultured in every 2 days. Old medium was removed from the flask, and cells were washed with PBS. The semi-adherent cells were lifted using accutase for 5 minutes at 37°C. After quenching with PBS, the cell suspension was centrifuged at 200 g for 4 minutes. The supernatant was removed, and the cell pellet resuspended in fresh media. The fresh cell suspension was transferred into a new flask with fresh media at the appropriate splitting ratio (1:5, 1:10) and cultured again in DMEM Glutamax with 5% FBS and P/S. Then BV-2 cells were plated onto 96-well poly-D-lysine coated clear flat-bottom plates (Perkin Elmer, Cat. No. 6055500) and left for 48 h. These cells were then treated with Aβ (co-)aggregates for 24 hours. After 24 hours, the medium was collected, snap-frozen in liquid nitrogen, and stored at -80°C for further TNF-α measurement. BV-2 cells were then fixed using 4% paraformaldehyde (PFA) for 20 minutes and permeabilized using 0.2% Triton X-100 for 15 minutes at room temperature. These BV-2 microglia cells were then washed twice with PBS and blocked for 1 hour at room temperature in blocking solution (3% horse serum in PBS containing 0.05% Triton X). Then 50 nM Alexa-Fluor-488-labeled EPR19392 and 25 nM Alexa-Fluor-647-labeled 6E10 were prepared in blocking solution and applied to cells and stored overnight at 4°C. Cells were washed with 3x PBS and stored in PBS until imaging. Fixed cells were imaged using the Opera Phenix high content imaging system (Perkin Elmer) using a 40x water objective (NA 1.1).

**Astrocyte Culture and immunohistochemistry:** Primary human astrocytes were purchased from ScienCell™ Research Laboratories (Carlsbad, CA, US). Astrocytes were cultured at 37°C in 5% $CO_2$ and 95% air, in phenol red-free Human Astrocyte Media, (ScienCell™, 1801prf) supplemented with fetal bovine serum (FBS) (ScienCell™,at 0025), penicillin/streptomycin (ScienCell™, Cat. No. 0503) and astrocyte growth supplement (ScienCell™, Cat. No. 1852). Cells were passaged when approaching ~85% confluency and were only used up to passage 7. Astrocytes were plated at appropriate densities on black bottom 96-well plates (Perkin Elmer, Cat. No. 6055500) and cultured for at least 24 hours prior to commencing cell treatments. Astrocytes were treated with Aβ (co-)aggregates for 24 hours. After 24 hours, the medium was collected, snap-frozen in liquid nitrogen, and stored at -80°C for further analysis. The cells were washed with 1xPBS and fixed using 4% PFA for 10 minutes at room temperature. Following fixation, cells were washed three times with 1x PBS and permeabilized in 0.3% triton-X 100 (Sigma, Cat. No. No. T8787) in PBS for 5 minutes at ambient temperature. Cells were blocked for 1 hour in 3% BSA-PBS and after that washed three times with PBS and incubated overnight with a mixture of 50 nM Alexa-Fluor-488-labeled

EPR19392 and 25 nM Alexa-Fluor-647-labeled 6E10 in PBS with 1% BSA and 0.1 mg/mL horse serum. These astrocytes were then rewashed three times with PBS and imaged in epifluorescence mode using the Opera Phenix High Content Imaging System (PerkinElmer®).

**Data analysis of cellular uptake assay:** Fifteen fields of view were imaged per well, with 3 or 4 wells per condition. For each field of view, images were taken in three channels: white differential image contrast (DIC) illumination to visualize the cells, 488 nm excitation (detecting apolipoprotein E antibody, Alexa-Fluor-488-labeled EPR19392) and 635 nm excitation (for Aβ42 antibody, Alexa-Fluor-647-labeled 6E10). First, DIC images were used to define cell perimeters, using a mask created around the border of the cells using an automated python-based program. The aim is to distinguish cell and non-cell areas based on the white light images, then apply the mask to the other two channels and measure the respective fluorescence intensities. Firstly, in order to automate the classification process, a model was applied to the program to learn how to distinguish the cell and non-cell areas. Ten white light images were manually labeled using the Trainable Weka Segmentation[6] in the Fiji interface[5] and three different class labels were created: background, cell area, and cluster area (where the cells were clustered) to generate a classifier model using these labeled images. A python code applied this model to all the white light images and generated probability maps for each class. Masks for cell areas were then created by only keeping the coordinates with a probability greater than 67% for the cell. Next, these masks were used in the other two fluorescence channels to extract the respective fluorescence intensities from the other two channels within each cell.

**ELISA to measure TNF-α in cell media:** To determine TNF-α secretion by BV-2 microglial cells and human astrocytes, cell media were collected after incubation of cells with different aggregates over 24 h and stored at -80°C until analyzed. TNF-α in the cell media was measured using the Duoset® enzyme-linked immunosorbent assay (ELISA) development system (R&D Systems, Abingdon, Oxfordshire, UK), according to the manufacturer's protocol.

**Membrane permeabilization assay:** The membrane permeabilization assays were performed as previously described[7]. Briefly, vesicles (mean diameter of 200 nm) were prepared by extrusion and five freeze-thaw cycles of a 100:1 mixture of phospholipids 16:0–18:1PC (Avanti Lipids, Cat. No. 850457C) and biotinylated lipids 18:1–12:0 Biotin PC (Avanti Lipids, Cat. No. 860563C). The lipid mixture was hydrated in 100 µM Cal-520 dye (Stratech, Cat. No. 21141) dissolved in HEPES buffer (50 mM, pH 6.5). Non-incorporated Cal-520 dye was separated from the dye-filled vesicles using size-exclusion chromatography, using a SuperdexTM 200 Increase 10/300 GL column attached to an AKTA pure system (GE Life Sciences) with a flow rate of 0.5 mL/min. Dye-filled vesicles were immobilized on coverslips

(VWR International, 22x22 mm, 631-0122), which were first cleaned using an argon plasma cleaner (PDC-002, Harrick Plasma) for ~45 minutes to remove any organic and fluorescent impurities. Frame-seal incubation chambers (9x9mm, Biorad, SLF-0601) were then affixed to the coverslips, and 50 µL of a mixture of 100:1 PLL-g-PEG (20 kDa PLL grafted with 2 kDa PEG and 3.5 Lys units/PEG Chains, SuSoS AG) and PLL-g-PEG biotin (20 kDa PLL grafted with 2 kDa PEG and 50% 3.4 kDa PEG-Biotin, SuSoS AG) at ~1 mg/mL was added to the coverslips and incubated for ~30 min. The coverslips were then washed three times with HEPES, and 50 µL of 0.1 mg/mL NeutrAvidin (Thermo Fisher, Cat. No. 31000) were added and incubated for 15 min. The coverslips were then rewashed three times with HEPES buffer, and 50 µL of the purified vesicles were added to the coverslips. Before imaging, 50 µL of $Ca^{2+}$-containing buffer (phenol red-free Leibovitz's L-15 Medium, Thermo Fisher Cat No. 21083027) was added, and blank images ($F_{blank}$) were recorded. The immobilized vesicles were then incubated with 50 µL of the aggregation mixture (100 nM in monomer equivalents) for 20 min and then reimaged ($F_{sample}$). Finally, 1 mg/mL of ionomycin (Cambridge bioscience, Cat. No. 1565-5) - which leads to $Ca^{2+}$ saturation inside the vesicles - was added, and the same areas were reimaged ($F_{ionomycin}$). The relative influx of $Ca^{2+}$ into an individual vesicle due to protein aggregates was then determined using the following equation: $Ca^{2+}$ influx = $F_{sample}$ - $F_{blank}$/$F_{ionomycin}$ - $F_{blank}$. The average degree of $Ca^{2+}$ influx was calculated by averaging the $Ca^{2+}$ influx in >200 individual vesicles. For the imaging, the Cal-520 dye was excited at 488 nm, and the emission from the dye was passed through a combination of a long-pass filter (BLP01-488R-25, Laser 2000) and a bandpass filter (FF01-520/44-25, Laser 2000) before being imaged using a Photometrics Evolve EMCCD camera. Images were acquired with a ~10 $W/cm^2$ power density with a scan speed of 20 Hz and bit depth of 16 bits.

**LDH cytotoxicity assay:** The assay was performed using an LDH Cytotoxicity Assay Kit (Thermo Fisher, Cat. No. C20303) using human neuroblastoma SH-SY5Y (ATCC) cells cultured in DMEM media without Phenol Red (Gibco, Thermo Fisher, Cat. No. 21063029) and supplemented with 10% Fetal Bovine Serum (Gibco, Thermo Fisher, Cat. No. A4766801). Cells were then treated with aliquots of the aggregation reactions at $t_1$ and $t_3$ time points (final Aβ42 concentration: 2 µM in monomer equivalents) for 12 hours, and the cell supernatant was assayed for LDH. The supernatant from cells treated with RIPA lysis buffer (Thermo Fisher, Cat. No. 89900) was used as a positive control, whereas media of the untreated SH-SY5Y cells were taken as a negative control. Next, the supplied reaction mixture (100 µL), which contains a substrate for the LDH activity detection reagent, was added to the medium according to the manufacturer's protocol. The reactions were quenched after 30 minutes with the Stop buffer (also supplied with the kit), and the absorbance at 480 nm was measured using a plate reader (Clariostar -BMG  Biotech).

**Immunoprecipitation assay:** Twenty-fiive microliters of Dynabeads Protein G (Invitrogen, Cat. No. 10007D) were mixed with 100 µL of a 100 nM solution of either HAE-4[8] or Mouse Isotype IgG2a kappa monoclonal [MG2a-53] (Biolegend) antibody in PBS and mixed in a tube rotator for 30 min at room temperature. The beads were then separated using a magnetic separation rack (DynaMag™-2 Magnet, Thermo Fisher, Cat. No. 12321D) and washed twice with PBS. Then, 300 µL of brain extract from *APOE4/4* homozygous AD patients or a mixture of *in vitro*-prepared lipidated and non-lipidated apoE-Aβ (150 µL each), from time-point $t_1$ were combined with the beads. This mixture was then rotated at 4°C for 3 h. After that, the magnetic rack was used to separate the beads and collect the supernatant for imaging and cellular assays.

**Statistical tests:** Data are displayed as the mean and error bars represent the standard deviation across replicates respectively. To assess the statistical significance of the differences between two populations we used unpaired two-sample t-tests. Multiple groups were compared using the one-way ANOVAs with post hoc Tukey-tests. Values of $p < 0.05$ were considered statistically significant. The number of independent biological replicates and technical replicates for each experiment as well as details of statistical tests for each experiment can be found in figure legends. Statistical tests were calculated with Origin 9.0 (OriginLab).

### References


1. Verghese, P.B., Castellano, J.M., Garai, K., Wang, Y., Jiang, H., Shah, A., Bu, G., Frieden, C., and Holtzman, D.M. (2013). ApoE influences amyloid-β (Aβ) clearance despite minimal apoE/Aβ association in physiological conditions. Proceedings of the National Academy of Sciences *110*, 1807–1816. 10.1073/pnas.1220484110.

2. Hong, W., Wang, Z., Liu, W., O'Malley, T.T., Jin, M., Willem, M., Haass, C., Frosch, M.P., and Walsh, D.M. (2018). Diffusible, highly bioactive oligomers represent a critical minority of soluble Aβ in Alzheimer's disease brain. Acta Neuropathologica *136*, 19–40. 10.1007/s00401-018-1846-7.

3. Di Antonio, M., Ponjavic, A., Radzevičius, A., Ranasinghe, R.T., Catalano, M., Zhang, X., Shen, J., Needham, L.-M., Lee, S.F., Klenerman, D., et al. (2020). Single-molecule visualization of DNA G-quadruplex formation in live cells. Nature Chemistry *12*, 832–837. 10.1038/s41557-020-0506-4.

4. Edelstein, A.D., Tsuchida, M.A., Amodaj, N., Pinkard, H., Vale, R.D., and Stuurman, N. (2014). Advanced methods of microscope control using μManager software. Journal of Biological Methods *1*, 1–10. 10.14440/jbm.2014.36.

5. Schindelin, J., Arganda-Carreras, I., Frise, E., Kaynig, V., Longair, M., Pietzsch, T., Preibisch, S., Rueden, C., Saalfeld, S., Schmid, B., et al. (2012). Fiji: an open-source platform for biological-image analysis. Nature Methods *9*, 676–682. 10.1038/nmeth.2019.

6. Arganda-Carreras, I., Kaynig, V., Rueden, C., Eliceiri, K.W., Schindelin, J., Cardona, A., and Sebastian Seung, H. (2017). Trainable Weka Segmentation: a machine learning tool for microscopy pixel classification. Bioinformatics *33*, 2424–2426. 10.1093/bioinformatics/btx180.

7. Flagmeier, P., De, S., Wirthensohn, D.C., Lee, S.F., Vincke, C., Muyldermans, S., Knowles, T.P.J.J., Gandhi, S., Dobson, C.M., and Klenerman, D. (2017). Ultrasensitive measurement of Ca2+ influx into lipid vesicles induced by protein aggregates. Angewandte Chemie International Edition *56*, 7750–7754. 10.1002/anie.201700966.

8. Liao, F., Li, A., Xiong, M., Bien-Ly, N., Jiang, H., Zhang, Y., Finn, M.B., Hoyle, R., Keyser, J., Lefton, K.B., et al. (2018). Targeting of nonlipidated, aggregated apoE with antibodies inhibits amyloid accumulation. The Journal of Clinical Investigation *128*, 2144–2155. 10.1172/JCI96429.

**Supplementary Information**

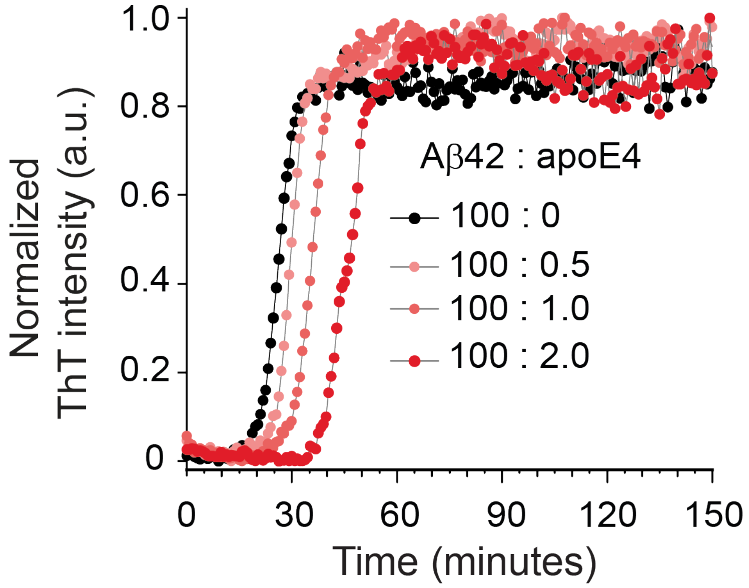


**Supplementary Figure 1.** Dependence of Aβ42 aggregation (4 μM) on the concentration of non-lipidated ApoE4 (0, 20, 40 and 80 nM), monitored by ThT fluorescence.

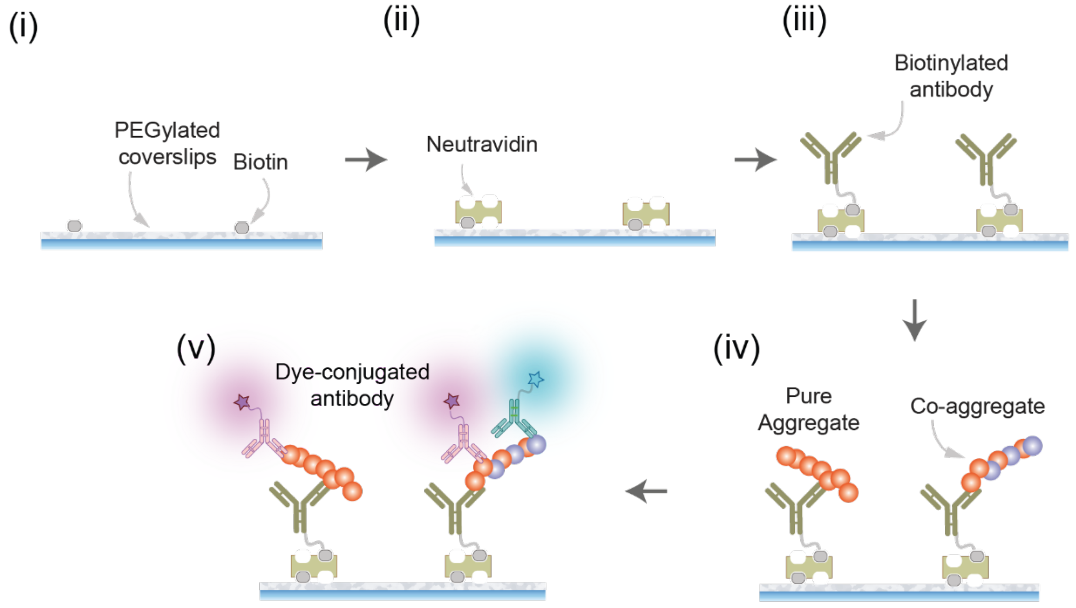


**Supplementary Figure 2.** Schematic of SiMPull assay.

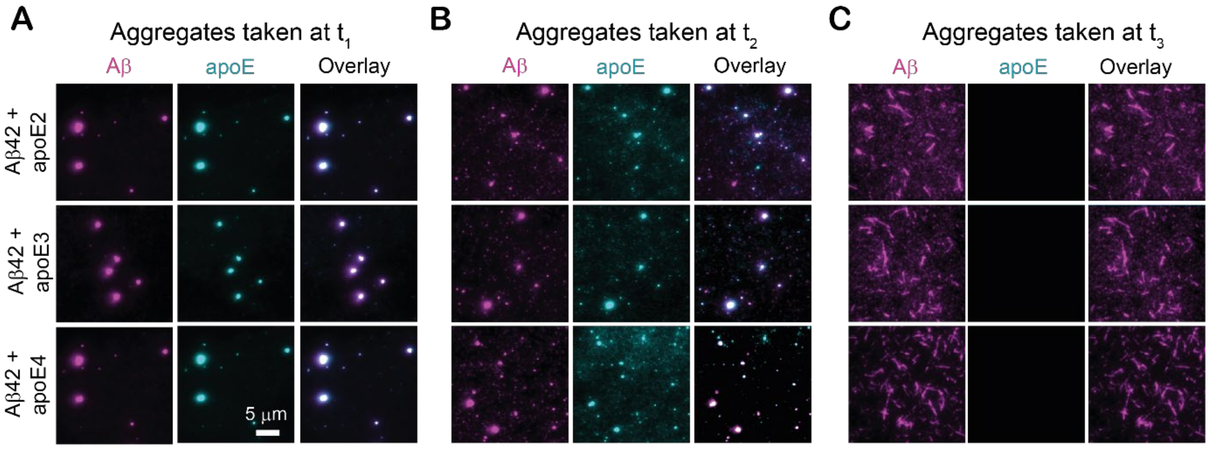


**Supplementary Figure 3. Colocalization of apoE with Aβ42 aggregates does not depend on the anti-apoE antibody used.** Aliquots of non-lipidated apoE-Aβ42 co-aggregation mixtures (aggregated at 4 μM Aβ42 and 80 nM apoE, then diluted 4-fold prior to pulldown) at $t_1$, $t_2$ and $t_3$ timepoints (as defined in Figure 1B) imaged using SiMPull with biotinylated 6E10 antibody for capture, and Aβ-specific Alexa-Fluor-647-labeled 6E10 (500 pM) and apoE-specific Alexa-Fluor-488-labeled F-9 (1 nM) as imaging antibodies. As in Figure 1 - where a different apoE antibody is used - apoE colocalizes with soluble Aβ42 aggregates, but not fibrillar aggregates.

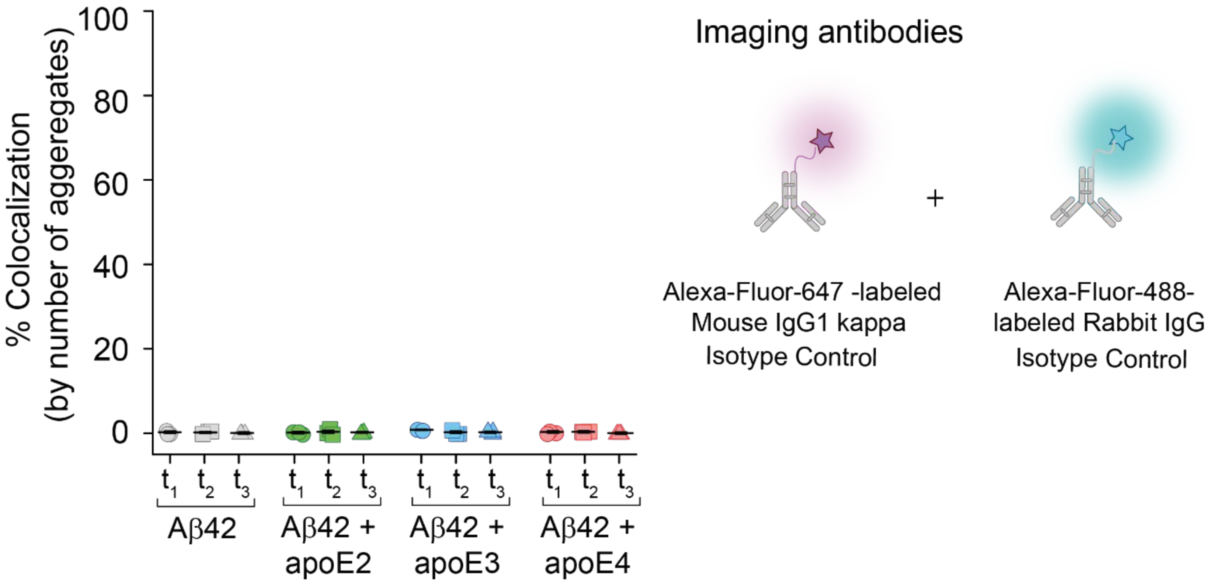


**Supplementary Figure 4. Isotype control antibodies generate no colocalization in two-color SiMPull imaging**. Aβ42 aggregates and non-lipidated apoE-Aβ42 co-aggregates formed at different stages of aggregation (defined in Figure 1B) and imaged with SiMPull using two isotype control antibodies. For all samples, no significant colocalization is observed.

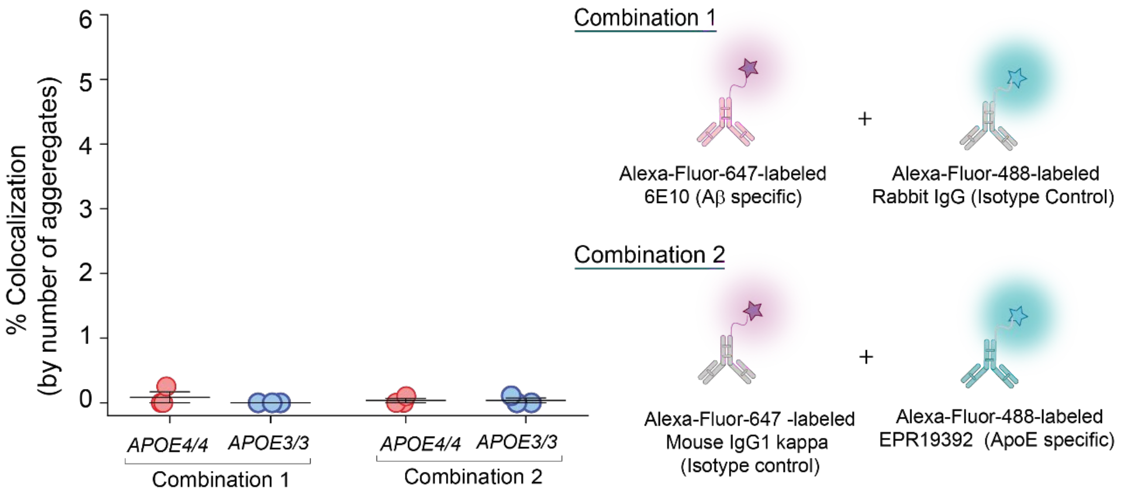


**Supplementary Figure 5. Aβ and apoE antibodies bind co-aggregates *via* their specific paratopes.** Aβ-containing aggregates captured from frontal cortex extracts of homozygous *APOE4* and *APOE3* AD patients imaged using two different combinations of imaging antibodies (n=3). In each experiment, one isotype control antibody is combined with an Aβ- or apoE-specific antibody respectively. No significant colocalization is observed, indicating that antibody binding is paratope mediated.

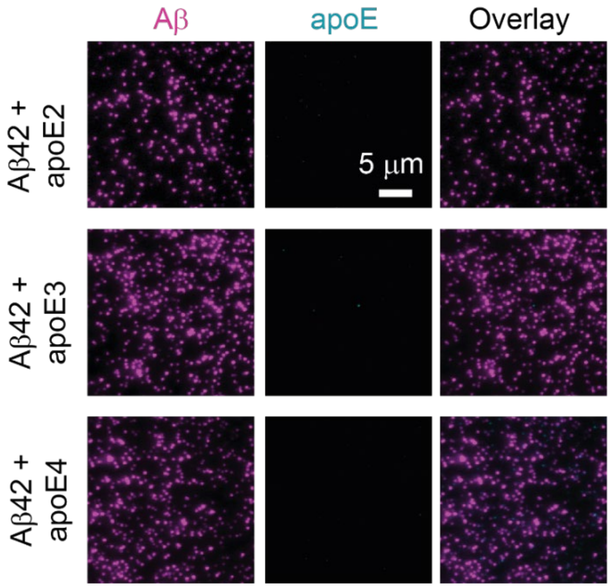


**Supplementary Figure 6. ApoE does not colocalize with pre-formed Aβ42 aggregates.** Aβ42 aggregates (4 μM monomer equivalents) taken at the end of the lag phase ($t_1$, as defined in Figure 1B), incubated in an ice bath with non-lipidated apoE (80 nM), diluted 4-fold in PBS then imaged using SiMPull.

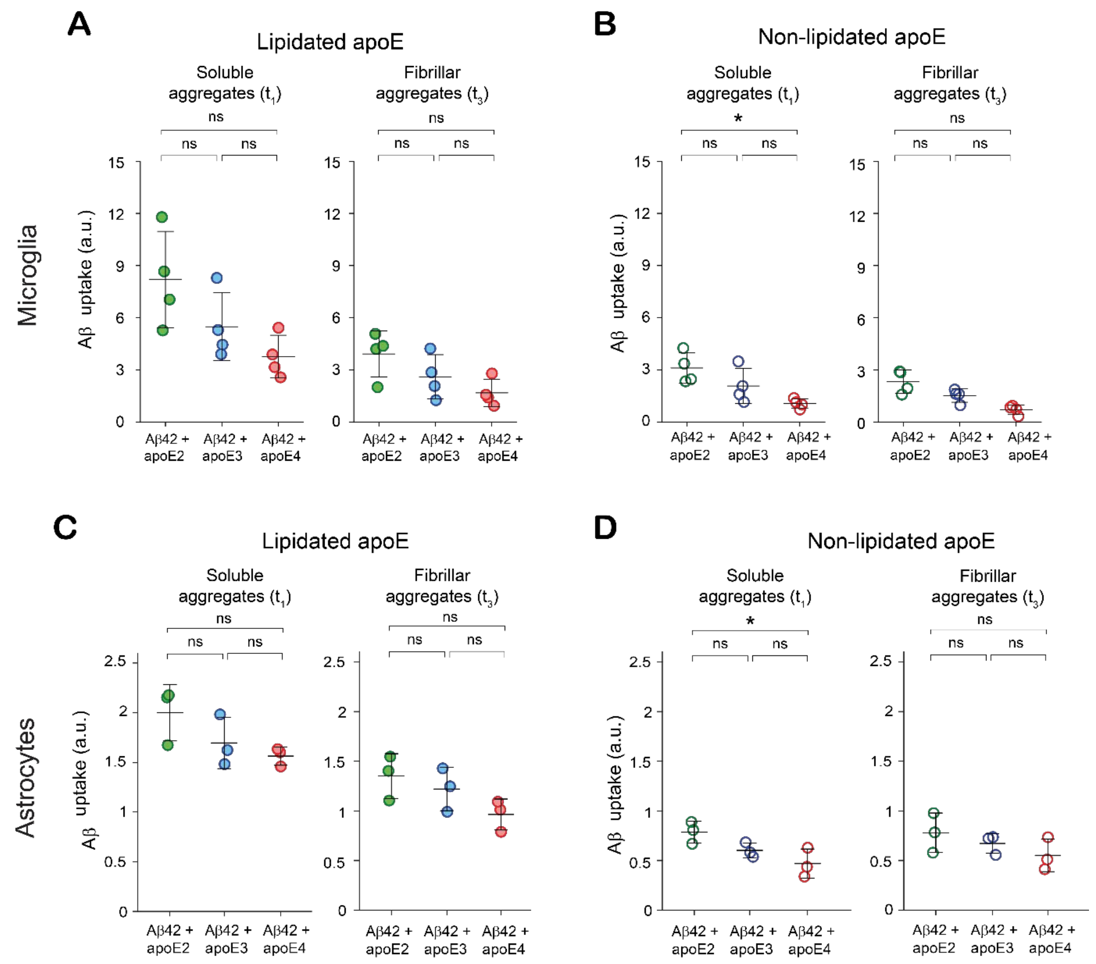


**Supplementary Figure 7. Uptake of apoE-Aβ co-aggregates by BV2 microglial cells and human astrocytes depends on apoE isoform.** Uptake of soluble ($t_1$) and fibrillar ($t_3$) Aβ aggregates by BV-2 microglia **(A, B)** and human astrocytes **(C, D)** formed in the presence of lipidated **(A)** and non-lipidated apoE **(B).** Units of uptake = integrated fluorescence of sample divided by the integrated fluorescence of internalized soluble Aβ42 aggregates at $t_1$ for each replicate (as in Figure 3). Data were averaged over four and three biological replicates for BV-2 microglia and astrocytes respectively; data are shown as individual data points and error bars represent standard deviation. Statistical significance was calculated using a one-way ANOVA with post-hoc Tukey. *$P < 0.05$, **$P < 0.01$, ***$P < 0.001$, ns, non-significant ($P > 0.05$).

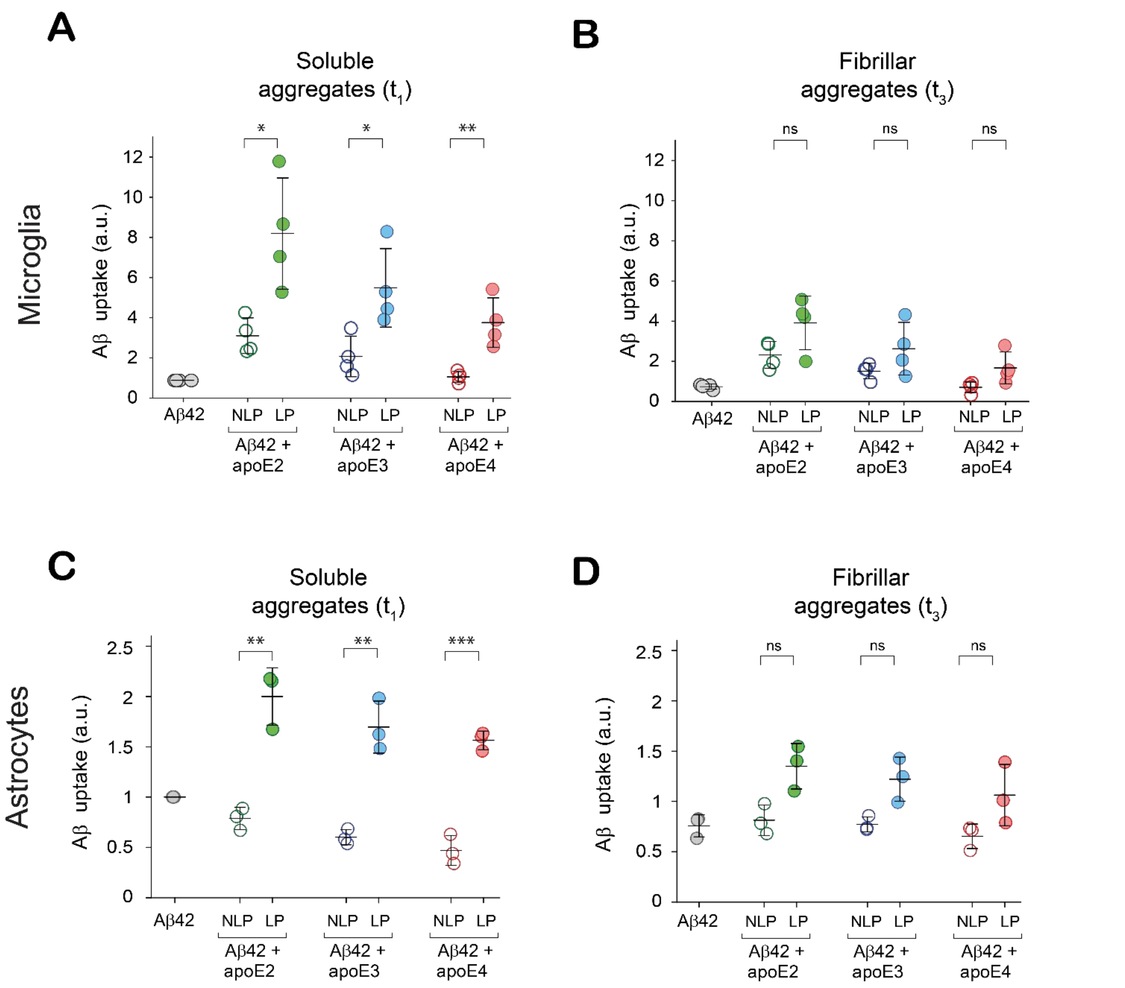


**Supplementary Figure 8. Association with lipidated apoE enhances uptake of Aβ aggregates by BV2 microglial cells and human astrocytes.** Uptake of Aβ aggregates by BV-2 microglia **(A,B)** and human astrocytes **(C,D)** formed in the presence of lipidated and non-lipidated apoE. Units of uptake = integrated fluorescence of sample divided by the integrated fluorescence of internalized soluble Aβ42 aggregates at $t_1$ for each replicate (as in Figure 3). Data points represent one of four (microglia) or three (astrocytes) biological replicates; error bars represent standard deviation. Statistical significance was calculated using an unpaired two sample T-test. *$P < 0.05$, **$P < 0.01$, ***$P < 0.001$, ns, non-significant ($P > 0.05$).

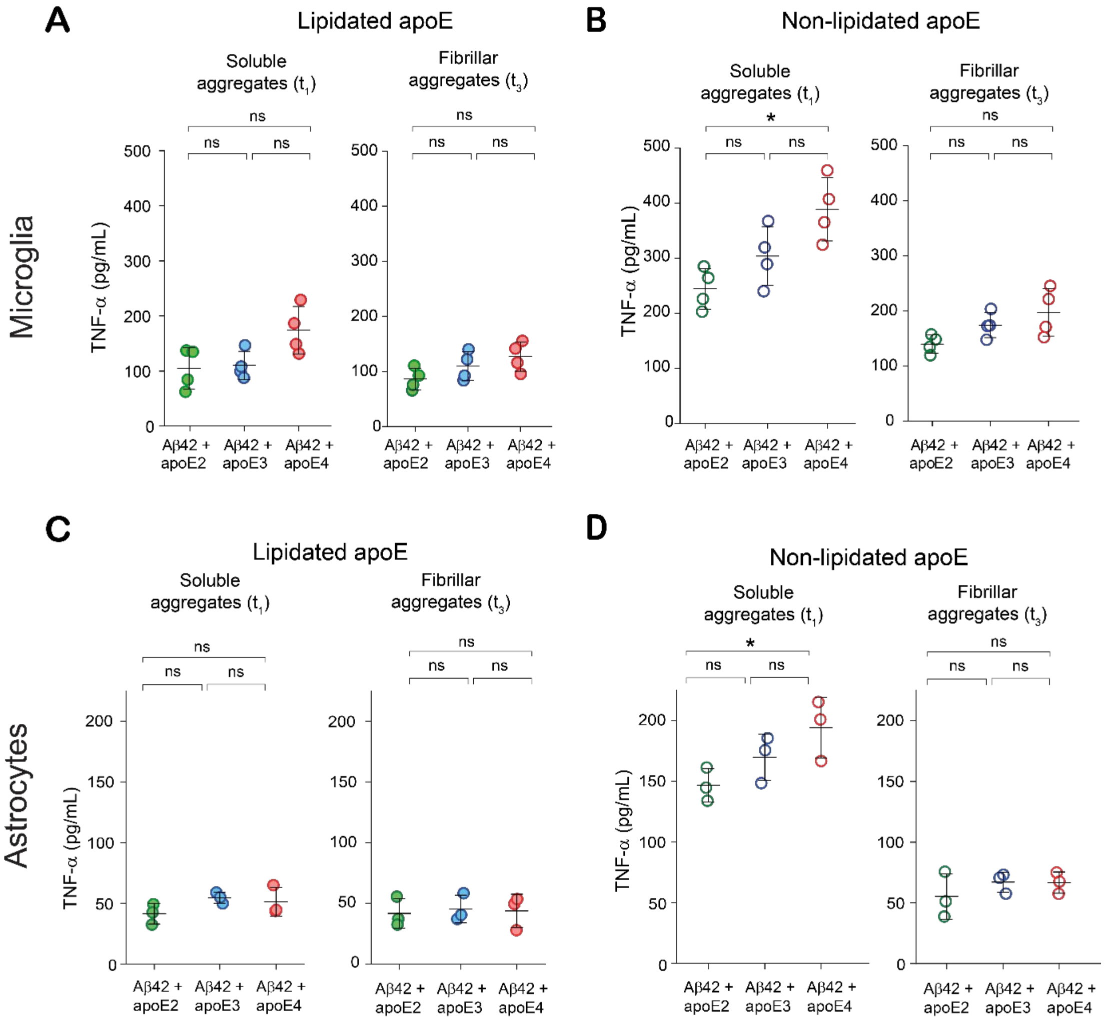


**Supplementary Figure 9. ApoE-Aβ co-aggregates inflame microglia and astrocytes in an isoform-dependent way.** TNF-α release by BV-2 microglia **(A-B)** and astrocytes **(C, D)** induced by soluble ($t_1$) and fibrillar ($t_3$) aggregates. Each data point represents one of four (for BV-2 cells) or three (for astrocytes) biological replicates; error bars represent standard deviation. Statistical significance was calculated using one-way ANOVA with post-hoc Tukey. *P < 0.05, **P < 0.01, ***P < 0.001, ns, non-significant (P > 0.05).

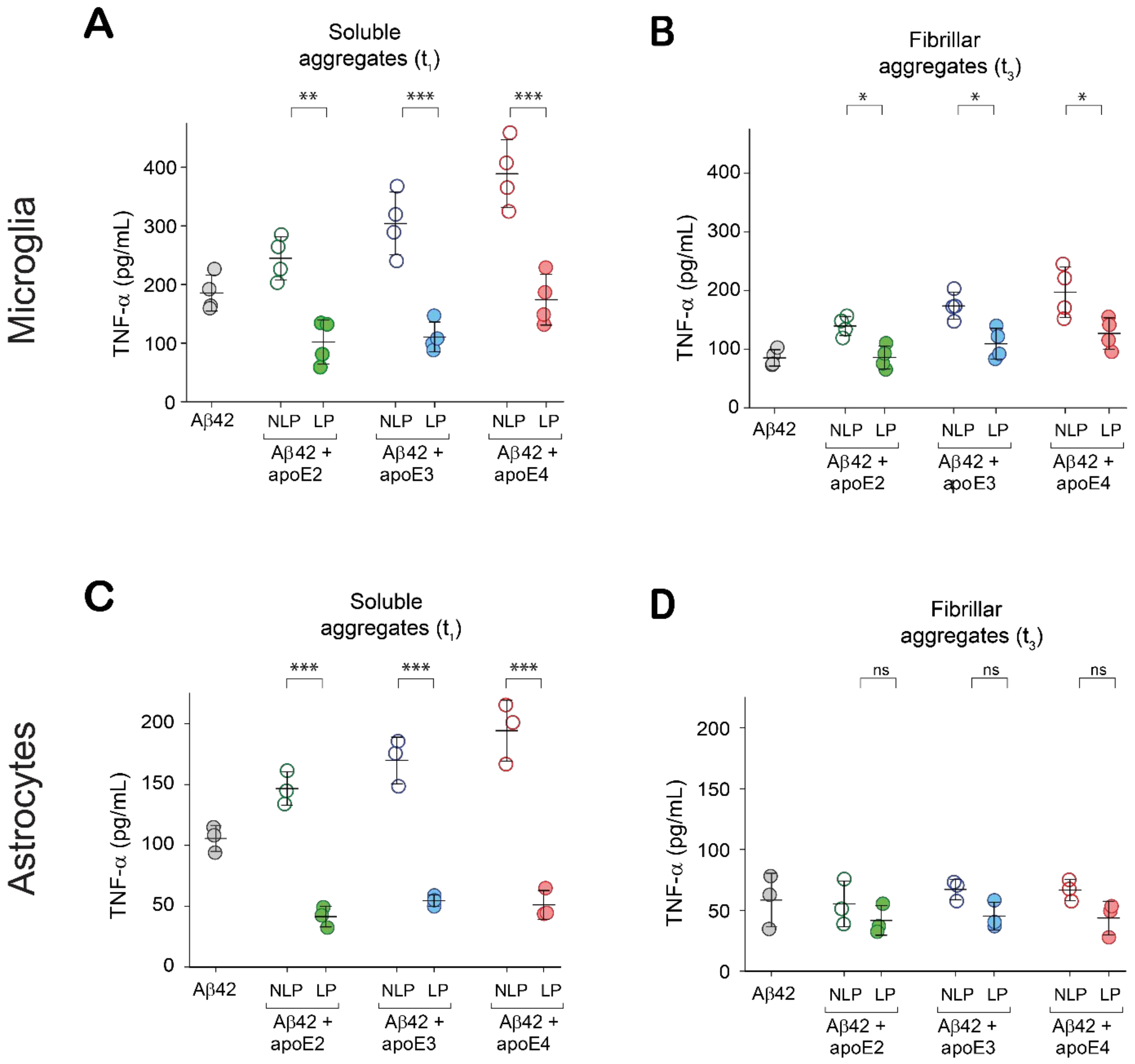


**Supplementary Figure 10. ApoE-Aβ co-aggregates inflame microglia and astrocytes in an apoE-lipidation-dependent manner.** TNF-α release by BV-2 microglia **(A-B)** and astrocytes **(C, D)** induced by soluble ($t_1$) and fibrillar ($t_3$) aggregates. Each data point represents one of four (for BV-2 cells) or three (for astrocytes) biological replicates; error bars represent standard deviation. Statistical significance was calculated using a two-sample t-test. *P < 0.05, **P < 0.01, ***P < 0.001, ns, non-significant (P > 0.05).

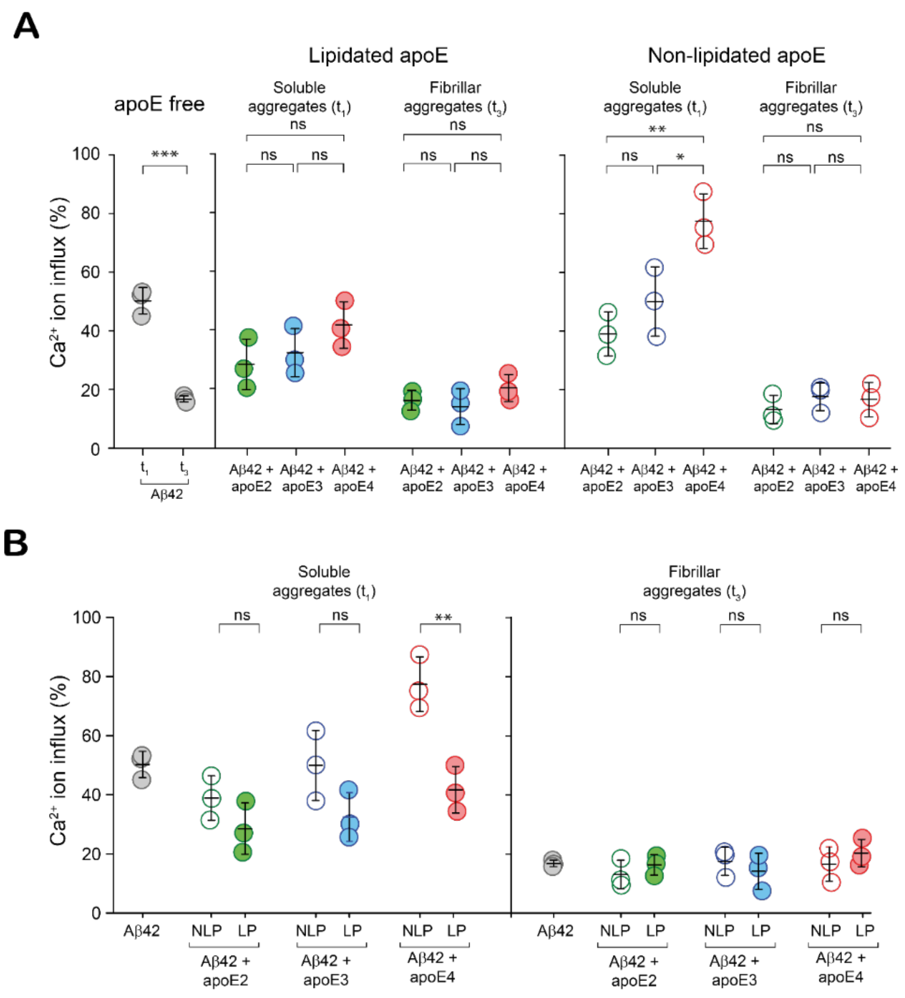


**Supplementary Figure 11. Isoforms and lipidation of apoE modulate the lipid membrane permeabilization ability of Aβ aggregates differently.** Comparison of lipid bilayer permeabilization by **(A)** apoE-Aβ co-aggregates containing different apoE isoforms and **(B)** lipidated and non-lipidated apoE-Aβ co-aggregates formed at $t_1$ (end of lag phase, where aggregates are mostly soluble co-aggregates) and $t_3$ (plateau phase, where aggregates are mostly Aβ-only fibrils) ([Aβ42] = 4 μM in monomer equivalents; [apoE] = 0 or 80 nM). $Ca^{2+}$ influx is referenced to the influx caused by the ionophore, ionomycin. Data points represent one of three technical replicates; error bars represent standard deviation and statistical significance was calculated using One-way ANOVA with post-hoc Tukey **(A)** and an unpaired two-sample t-test **(B)**. *$P < 0.05$, **$P < 0.01$, ***$P < 0.001$, NS, non-significant ($P > 0.05$).

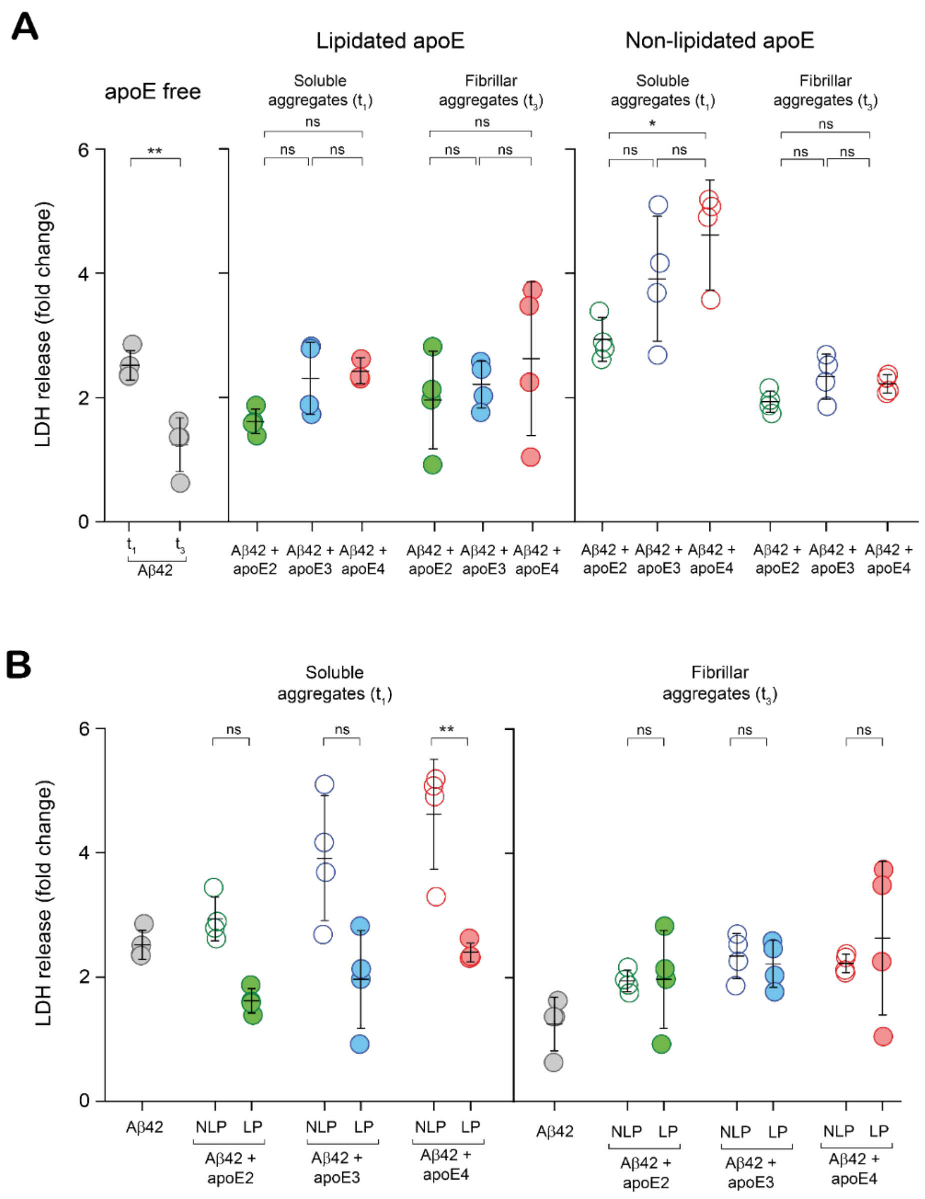


**Supplementary Figure 12. Isoforms and lipidation of apoE modulate the Aβ-aggregate-induced LDH release differently.** Comparison of LDH release elicited by **(A)** apoE-Aβ co-aggregates containing different apoE isoforms and **(B)** lipidated and non-lipidated apoE-Aβ co-aggregates formed at $t_1$ (end of lag phase, where aggregates are mostly soluble co-aggregates) and $t_3$ (plateau phase, where aggregates are mostly Aβ-only fibrils) ([Aβ42] = 4 μM in monomer equivalents; [apoE] = 0 or 80 nM). Units of LDH release = LDH release elicited by sample divided by LDH release elicited by buffe for each replicate. Data points represent one of four biological replicates; error bars represent standard deviation and statistical significance was calculated using One-way ANOVA with post-hoc Tukey **(A)** and an unpaired two-sample t-test **(B)**. *P < 0.05, **P < 0.01, ***P < 0.001, NS, non-significant (P > 0.05).
